# Long non-coding RNA *GRASLND* enhances chondrogenesis via suppression of interferon type II signaling pathway

**DOI:** 10.1101/650010

**Authors:** Nguyen P.T. Huynh, Catherine C. Gloss, Jeremiah Lorentz, Ruhang Tang, Jonathan M. Brunger, Audrey McAlinden, Bo Zhang, Farshid Guilak

## Abstract

Long non-coding RNAs (lncRNAs) play critical roles in regulating gene expression and cellular processes; however, their roles in musculoskeletal development, disease, and regeneration remain poorly understood. Here, we identified a novel lncRNA, Glycosaminoglycan Regulatory ASsociated Long Non-coDing RNA (*GRASLND*) as a regulator of mesenchymal stem cell (MSC) chondrogenesis, and we investigated its basic molecular mechanism and its potential application towards regenerative medicine. *GRASLND,* a primate-specific lncRNA, is upregulated during MSC chondrogenesis and appears to act directly downstream of SRY-Box 9 (SOX9), but not Transforming Growth Factor Beta 3 (TGF-β3). Utilizing the established model of pellet formation for MSC chondrogenesis, we showed that the silencing of *GRASLND* resulted in lower accumulation of cartilage-like extracellular matrix, while *GRASLND* overexpression, either via transgene ectopic expression or by endogenous activation via CRISPR, significantly enhanced cartilage matrix production. *GRASLND* acts to inhibit interferon gamma (IFN-γ) by binding to Eukaryotic Initiation Factor-2 Kinase EIF2AK2. We further demonstrated that *GRASLND* exhibits a protective effect in engineered cartilage against interferon type II across different sources of chondroprogenitor cells. Our results indicate an important role of *GRASLND* in regulating stem cell chondrogenesis, as well as its therapeutic potential in the treatment of cartilage-related diseases, such as osteoarthritis.

**Significance:** Long non-coding RNAs (lncRNAs) play critical roles in gene regulation and cellular physiology; however, the role of lncRNAs in controlling stem cell chondrogenesis remains to be determined. Here, we utilized next generation sequencing of adult stem cell chondrogenesis to identify a set of potential lncRNA candidates involved in this process. We identified lncRNA Glycosaminoglycan Regulatory ASsociated Long Non-coDing RNA (*GRASLND*) and characterized its molecular mechanism of action. We described a novel role of *GRASLND* in positive regulation of chondrogenesis via its inhibition of type II interferon. Importantly, we showed that overexpression of *GRASLND* augments stem cell chondrogenesis, providing a promising approach to enhancing stem cell chondrogenesis and cartilage regeneration.

## Introduction

Articular cartilage is an aneural, avascular tissue and has little or no capacity for intrinsic repair (1). While there are currently no effective treatments available for cartilage repair, focal cartilage or osteochondral lesions generally progress to osteoarthritis (OA), a progressive degenerative disease characterized by changes in the articular cartilage and remodeling of other joint tissues such as the synovium and subchondral bone. Thus, there remains an important need for regenerative therapies that can enhance cartilage repair through tissue engineering or cell therapy approaches (2-7).

In this regard, adult stem cells such as bone marrow derived mesenchymal stem cells (MSCs) or adipose-derived stem cells (ASCs) provide a readily accessible source of multipotent cells that show significant promise for regenerative medicine (8–11). When cultured in a defined cocktail supplemented with Transforming Growth Factor Beta 3 (TGF-β3), MSCs produce a cartilaginous matrix that is rich in glycosaminoglycan (GAG) and collagen type II (COLII) (12, 13). However, the complete pathway involved in MSC chondrogenesis is not fully deciphered, and a detailed understanding of the gene regulatory networks that control this process could provide new insights that accelerate and improve cartilage regeneration from endogenous or exogenously grafted MSCs.

Increasing evidence suggests that such gene regulatory pathways operational in stem cell differentiation may rely not only on protein-coding RNAs, but also on non-coding RNAs (ncRNAs). Non-coding RNAs (ncRNAs) were initially difficult to identify because they neither possessed open reading frames, nor were they evolutionarily highly conserved (14). In one of the first landmark studies, chromatin-state mapping was used to identify transcriptional units of functional large intervening non-coding RNAs (lincRNAs) that were actively transcribed in regions flanking protein-coding loci (15), and follow-up loss-of-function studies indicated that these lincRNAs were indeed crucial for the maintenance of pluripotency in embryonic stem cells (16). There is a growing understanding of long non-coding RNA (lncRNA) function in a multitude of tissues and cellular processes. For example, detailed mechanistic studies on the role of lncRNAs in X chromosome inactivation (17), or nervous system development and functions (18, 19) have been previously reported. However, knowledge of their roles in the musculoskeletal system, particularly in chondrogenesis remains limited. Only a handful of functional studies have been carried out in this regard. For example, lncRNA-HIT (HOXA Transcript Induced by TGFβ) (20) has been shown to play a role in epigenetic regulation during early limb development. Other studies have implicated a specific lncRNA, ROCR (Regulator of Chondrogenesis RNA) (21), to act upstream of SRY-Box 9 (SOX9) and regulate chondrocyte differentiation (22).

As one of their many modes of actions, lncRNAs are also known to regulate and modulate various signaling cascades involved in the control of gene regulatory networks. Therefore, there may exist a connection between lncRNA candidates and signaling pathways previously known to play a role in the musculoskeletal system development. More specifically, there is growing evidence for the role of interferon (IFN) in skeletal tissue development and homeostasis (23–31). There are two main types of IFN. Type I includes mainly IFN alpha (IFN-α) and IFN beta (IFN-β) that form complexes with Interferon Alpha and Beta Receptors (IFNARs), activating the Janus Kinase/ Signal Transducers and Activators of Transcription (JAK/STAT) pathway by phosphorylation of STAT1 (Signal Transducer and Activator of Transcription 1) and STAT2 (Signal Transducer and Activator of Transcription 2). Phosphorylated STAT1/STAT2 then form complexes with IRF9 (IFN Regulatory Factor 9) and translocate into the nucleus to activate downstream targets via the interferon-stimulated responsible element (ISRE) DNA binding motif. Type II, on the other hand, relies on activation of the JAK/STAT pathway following the binding of IFN gamma (IFN-γ) to Interferon Gamma Receptors (IFNGRs). This subsequently results in the phosphorylation and dimerization of STAT1 that translocates into the nucleus and induces downstream targets via the gamma activated sequence (GAS) DNA binding element (32–34).

Although interferons (IFN) are widely known for their antiviral response, they can also act in other aspects of cellular regulation (33). Interestingly, IFN-γ has been implicated in non-viral processes, most notably its priming effect in auto-immune diseases such as lupus nephritis, multiple sclerosis, or rheumatoid arthritis (35). An additional goal of this study was to elucidate the link between IFN-γ and our lncRNA candidate, and how this interaction could potentially play a role in MSC chondrogenesis and cartilage tissue engineering.

In a recent publication, we used high-depth RNA sequencing to map the transcriptomic trajectory of MSC chondrogenesis (36). This dataset provides a unique opportunity to identify candidate genes for subsequent functional characterization as regulators of chondrogenesis. Here, we used bioinformatic approaches to integrate our RNA-seq data with other publicly available datasets, applying a rational and systematic data mining method to define a manageable list of final candidates for follow-up experiments. As a result, we identified *RNF144A-AS1* to be a crucial regulator of chondrogenesis, and proposed the name Glycosaminoglycan Regulatory ASsociated Long Non-coDing RNA (*GRASLND*). We showed that *GRASLND* enhances chondrogenesis by acting to suppress the IFN-γ signaling pathway, and this effect was prevalent across different adult stem cell types and conditions. Together, these results highlight novel roles of *GRASLND* and its modulation of IFN in stem cell chondrogenesis, as well as its therapeutic potential to enhance cartilage regeneration.

## Results

### *RNF144A-AS1* is crucial to and specifically upregulated in chondrogenesis

First, we utilized our published database on MSC chondrogenesis (GSE109503) (36) to identify long non-coding RNA candidates. We investigated the expression patterns of MSC markers *(ALCAM, ENG, VCAM1)*, chondrogenic markers (*ACAN*, *COL2A1*, *COMP*), and SOX transcription factors (*SOX5, SOX6, SOX9*) (Figure S1A). Pearson correlation analysis revealed 141 lncRNAs whose expression was highly correlated to those of MSC markers, 40 lncRNAs to chondrogenic markers, and 17 lncRNAs to SOX transcription factors (Figure S1B, C). Among those, two were downregulated and two were upregulated upon ectopic SOX9 overexpression (Table 1) (GSE69110) (37). To validate their functions in chondrogenesis, we systematically designed small hairpin RNAs (shRNAs) targeting each candidate and assessed knockdown effect after 21 days of chondrogenic induction. We successfully designed two target shRNAs for each of three candidates, and one target shRNA for the other candidate (Figure S2). We showed that knockdown of two out of three MSC-related lncRNAs did not influence the production of glycosaminoglycans (GAG) - an important extracellular matrix component in cartilage (Figure S2). While these lncRNAs may have other regulatory functions in MSCs, their roles in chondrogenesis appeared to be minimal. Moreover, we found that lower levels of MSC-correlated lncRNAs did not prime the MSCs toward chondrogenesis. However, knockdown of *RNF144A-AS1* (*RNF144A Antisense RNA 1*) resulted in decreased expression of chondrogenic markers (*COL2A1*, *ACAN*), and upregulation of apoptotic (*CASP3*) and cellular senescence (*TP53*) markers (Figure 1A, B). This effect was not due to nonspecific cytotoxicity of examined shRNAs, as released levels of lactase dehydrogenase (LDH) were similar among control and shRNA-expressing cells (Figure S3). In addition, biochemical assays indicated a reduction in GAG deposition (p < 0.0001) as well as DNA and GAG/DNA levels (p < 0.001) (Figure 1 C-E). Histologically, we observed the same phenotypic loss of GAG and collagen type II in the extracellular matrices (ECM) of pellet samples with *RNF144A-AS1* targeted shRNAs, while the scrambled controls displayed explicit staining of these proteins (Figure 1F). Taken together, this data indicates that *RNF144A-AS1* may be required for both cellular proliferation and cartilage-like matrix production.

**Figure 1:**
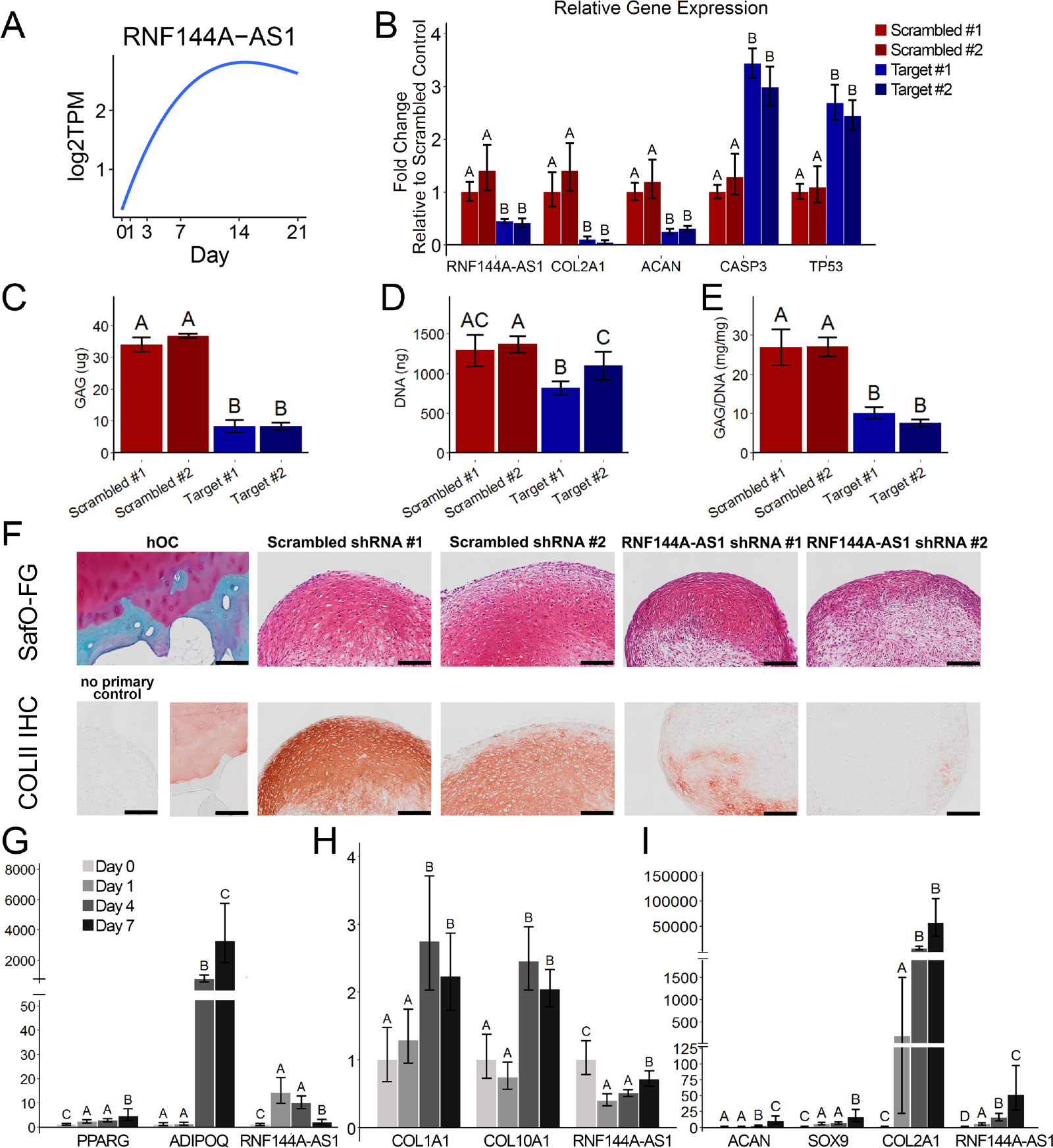
*RNF144A-AS1* is important and specifically upregulated in MSC chondrogenesis. (A) Expression pattern of *RNF144A-AS1* in chondrogenesis (GSE109503 (36)). Log2TPM: log transformed value of transcripts per million (TPM). (B) Effect of *RNF144A-AS1* knockdown on chondrogenic, apoptotic, and cell cycle inhibition markers (n = 5). (C-E) Effect of *RNF144A-AS1* knockdown on pellet matrix synthesis (n = 5). (F) Representative histological images of day 21 MSC pellets. Scale bar = 200 µm. *SafO-FG: SafraninO-Fast Green staining. COLII IHC: collagen type II immunohistochemistry. hOC: human osteochondral control.* (G-I) qRT-PCR analysis of MSC samples cultured in (G) adipogenic condition (n=6), (H) osteogenic condition (n=6), (I) chondrogenic condition (n=3-4). One-way ANOVA followed by Tukey post-hoc test (α=0.05). Groups of different letters are statistically different from one another.

**Table 1:** Long non-coding RNA candidates shortlist

| Gene symbol | Gene name | ENSEMBL gene ID | Relationship to MSC chondrogenesis | Relationship to SOX9 |
| --- | --- | --- | --- | --- |
| LOXL1-AS1 | LOXL1 antisense RNA 1 | ENSG00000261801 | Correlated with MSC markers' expression | Downregulated upon SOX9 overexpression |
| MIR4435-2HG<br>Gene synonym:<br>MIR4435-1HG | MIR4435-2 host gene | ENSG00000172965 | Correlated with MSC markers' expression | Downregulated upon SOX9 overexpression |
| HMGA2-AS1<br>Gene synonym:<br>RP11-366L20.2 | HMGA2 antisense RNA 1 | ENSG00000197301 | Correlated with MSC markers' expression | Upregulated upon SOX9 overexpression |
| RNF144A-AS1 | RNF144A antisense RNA 1 | ENSG00000228203 | Correlated with chondrogenic markers' expression | Upregulated upon SOX9 overexpression |

To establish whether *RNF144A-AS1* expression is specific to chondrogenesis or involved in other differentiation pathways, MSCs were induced towards adipogenic, osteogenic, or chondrogenic lineages, and *RNF144A-AS1* expression was measured at various timepoints throughout these processes. Successful differentiation was observed with an increase in lineage-specific markers: *PPARG* (*Peroxisome Proliferator Activated Receptor Gamma*) and *ADIPOQ* (*Adiponectin, C1Q And Collagen Domain Containing*) for adipogenesis, *COL1A1* (*Collagen Type I Alpha 1 Chain*) and *COL10A1* (*Collagen Type X Alpha Chain 1*) for osteogenesis, and *ACAN* (*Aggrecan*), *SOX9* (*SRY-Box 9*) and *COL2A1* (*Collagen Type II Alpha Chain 1*) for chondrogenesis (Figure 1 G-I). We found that *RNF144A-AS1* expression was particularly enriched as chondrogenesis progressed (Figure 1I). In contrast, *RNF144A-AS1* peaked at earlier timepoints during adipogenesis but decreased at later time points (Figure 1G), and downregulated when MSCs underwent osteogenic induction (Figure 1H), indicating that *RNF144A-AS1* is specifically upregulated in chondrogenesis. Furthermore, we speculate that *RNF144A-AS1* may display inhibitory effects on osteogenesis and adipogenesis, thus being downregulated during these processes.

To validate these gene expression findings, we performed RNA fluorescence *in situ* hybridization (FISH) throughout the time course of MSC chondrogenesis. Pellets exhibited *RNF144A-AS1* FISH signals at later time points during chondrogenic differentiation, consistent with RNA-seq data (Figure 2A). Next, to confirm *RNF144A-AS1* subcellular location, we performed qRT-PCR on isolated nuclear and cytoplasmic fractions of day 21 MSC pellets (Figure 2B). We compared the subcellular expression patterns of *RNF144A-AS1* to *NEAT1* (*Nuclear Paraspeckle Assembly Transcript 1*) and *GAPDH* (*Glyceraldehyde 3-Phosphate Dehydrogenase*). *NEAT1* is a lncRNA previously characterized to localize to the nucleus (38, 39), and *GAPDH* is an mRNA and thus should be exported to the cytoplasm for protein synthesis. Consistent with previous findings, *NEAT1* displayed lower expression in the cytoplasmic compared to the nuclear fraction, in contrast to *GAPDH*. By this measurement, *RNF144A-AS1* exhibited higher expression in the cytoplasm, indicating its cytoplasmic subcellular location. Our finding was recapitulated by RNA *in situ* hybridization followed by confocal microscopy (Figure 2C). Interestingly, since *RNF144A-AS1* showed punctate labeling, we speculate that this lncRNA may function in the form of an RNA-protein complex.

**Figure 2:**
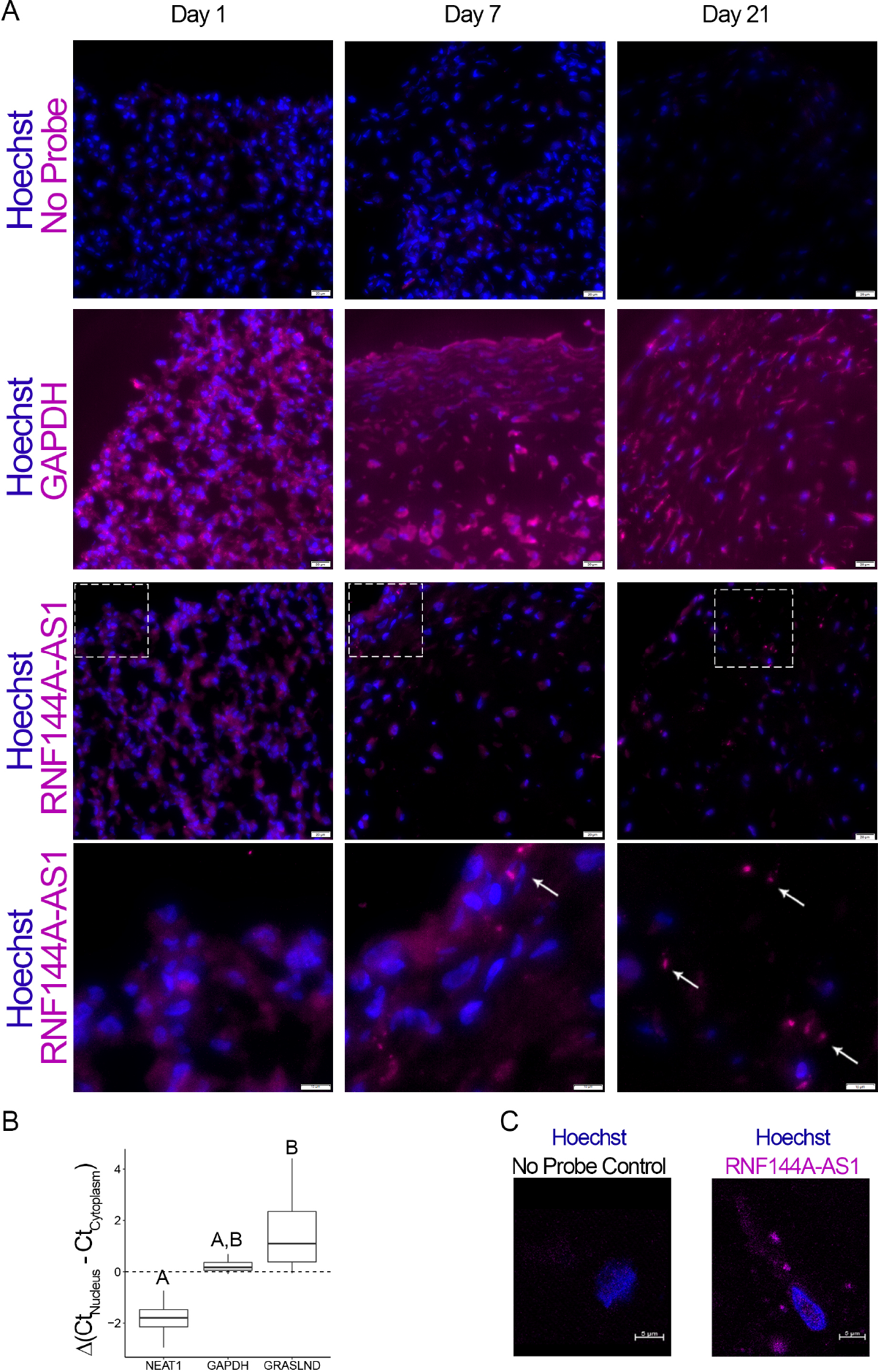
*RNF144A-AS1* is localized to the cytoplasm. (A) RNA in situ hybridization of MSC-derived pellets at different time points during chondrogenesis. *GAPDH* and *RNF144A-AS1* probes were hybridized on separate slides. Scale bar = 20 µm. (B) qRT-PCR of nuclear versus cytoplasmic fraction of day 21 MSC pellets (n=4). (C) Confocal microscopy on MSC-derived pellets. Scale bar = 5 µm. One-way ANOVA followed by Tukey post-hoc test (*α*=0.05). Groups of different letters are statistically different from one another.

### Characterization of *RNF144A-AS1*

We examined the characteristics of *RNF144A-AS1* by first exploring its evolutionary conservation. Except for exon 1, the genomic region of *RNF144A-AS1* is highly conserved in primates (*Homo sapiens, Pan troglodytes,* and *Rhesus macaque*) whose common ancestor traced back to 25 million years ago (40), while sequences are less conserved in other mammals (Figure 3A). This suggests that *RNF144A-AS1* may belong to a group of previously identified primate-specific lncRNAs (41, 42).

**Figure 3:**
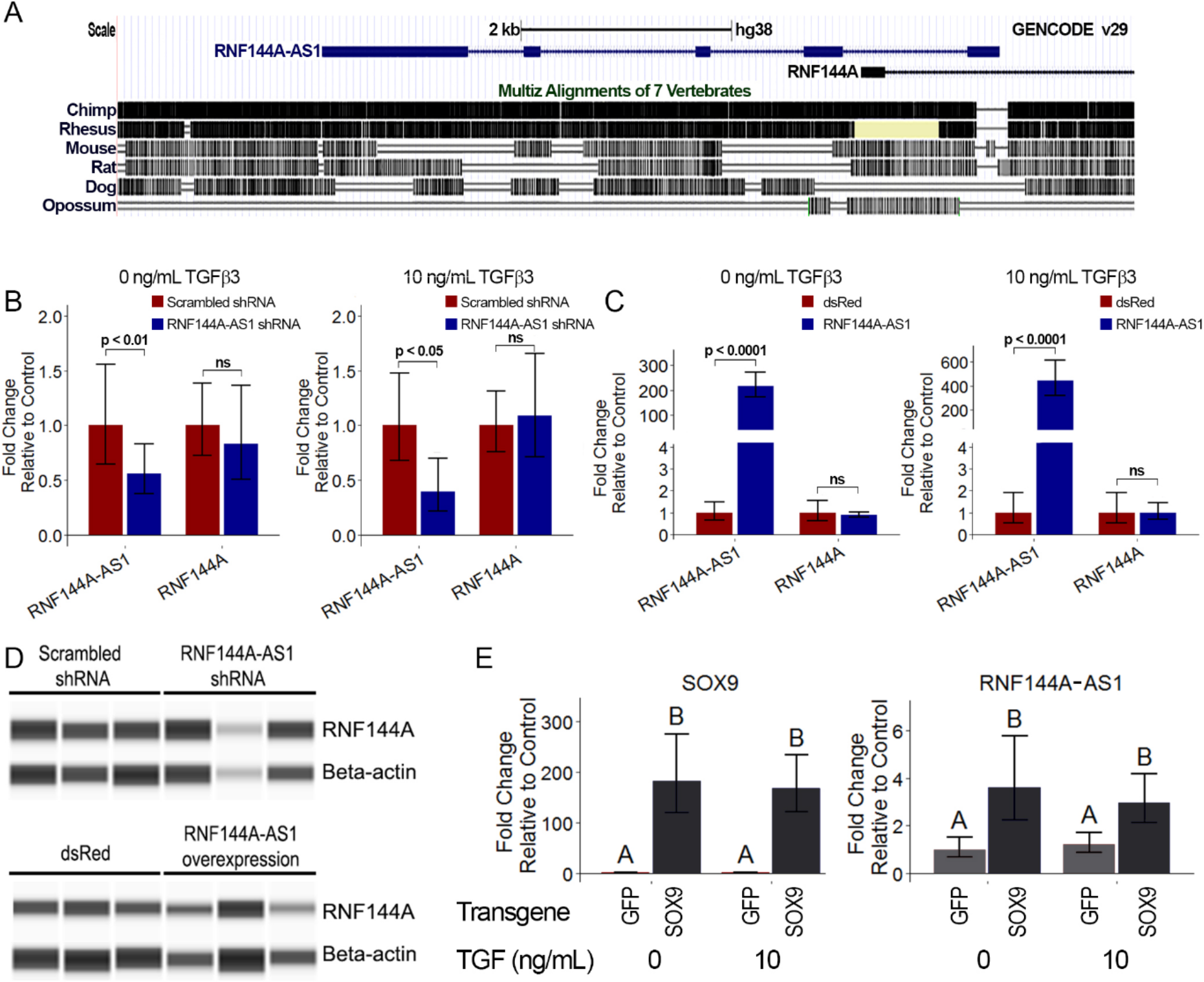
*RNF144-AS1* relationship to RNF144A and SOX9. (A) *RNF144A-AS1* genomic location and conservation across different species. Data retrieved from UCSC Genome Browser. (B) Knockdown of *RNF144A-AS1* and expression of *RNF144A* (n=4). (C) Overexpression *RNF144A-AS1* and expression of *RNF144A* (n=4). Welch’s t-test. (D) Protein amount of RNF144A by western blot in variation of *RNF144A-AS1* levels. Lanes indicate biological replicates. (E) *RNF144A-AS1* level in GFP- or SOX9-transduced MSCs under different doses of TGF-β3 (n=6). Two-way ANOVA followed by Tukey post-hoc test (α=0.05) on the effect of SOX9 overexpression (p < 0.0001) and doses of TGF-β3 (p > 0.05). The interaction between two tested factors (SOX9 overexpression and TGF-β3 doses) was not significant (p > 0.05). Groups of different letters are statistically different. *ns: not significant*.

Per GENCODE categorization, the AS (antisense) suffix indicates a group of lncRNAs that are positioned on the opposite strand, with overlapping sequences to their juxtaposed protein-coding genes. Often, these lncRNAs play a role in regulating the expression of their protein-coding counterparts (22). Therefore, we set out to examine whether this is also the case for *RNF144A-AS1* (Figure 3B-C). Neither knockdown nor overexpression of *RNF144A-AS1* affected RNF144A transcript levels in MSCs cultured with or without TGF-*β*3. Moreover, RNF144A protein translation also remained unaffected with variations of *RNF144A-AS1* levels, as indicated by western blot (Figure 3D). These results indicate that *RNF144A-AS1* is not involved in the regulation of RNF144A. For these reasons, we proposed an alternative name for *RNF144A-AS1*: *GRASLND* - *Glycosaminoglycan Regulatory ASsociated Long Non-coDing RNA*.

Next, we explored the signaling axis of *GRASLND*. Data mining and computational analysis on earlier published data suggested that *GRASLND* was a downstream effector of SOX9 (GSE69110) (37). When SOX9 was overexpressed in fibroblasts, *GRASLND* expression was increased (∼ 2-fold). We further confirmed this by utilizing SOX9 transgene overexpression in our MSCs culture (Figure 3E). Interestingly, while TGF-β3 has been demonstrated to act upstream of SOX9, exogenous addition of this growth factor alone did not result in enhanced *GRASLND* expression. It is notable that SOX9 levels in GFP controls were indistinguishable between TGF-β3 conditions at the time of investigation (1 week in monolayer culture), consistent with our previous finding that SOX9 was not upregulated until later timepoints in MSC chondrogenesis (36). Therefore, TGF-β3, despite being a potent growth factor, is not sufficient to elevate *GRASLND* expression. Instead, *GRASLND* appeared to be a downstream target of SOX9.

### Enhanced chondrogenesis for cartilage tissue engineering with *GRASLND*

As knockdown of *GRASLND* inhibited GAG and collagen deposition, we sought to investigate whether overexpression of *GRASLND* would enhance chondrogenesis. We assessed this question by both transgene ectopic expression and by CRISPR-dCas9 (Clustered regularly interspaced short palindromic repeats – catalytically dead Cas9) mediated in-locus activation.

We designed our lentiviral transfer vector to carry a BGH-pA (Bovine Growth Hormone Polyadenylation) termination signal downstream of *GRASLND* to allow for its correct processing (Figure S4A). Additionally, *GRASLND* was also driven under a doxycycline inducible promoter, enabling the temporal control of its expression. We utilized this feature to induce *GRASLND* only during chondrogenic culture (Figure 4A). This experimental design focused solely on the role of *GRASLND* during chondrogenesis, while successfully eliminating its effect in MSC maintenance and expansion from our analysis. As control, a vector encoding the Discosoma sp. red fluorescent protein (dsRed) coding sequence in place of *GRASLND* was utilized. Since doxycycline was most potent at 1 µg/mL (Figure S4 B-C), this dose was used for all following experiments.

**Figure 4:**
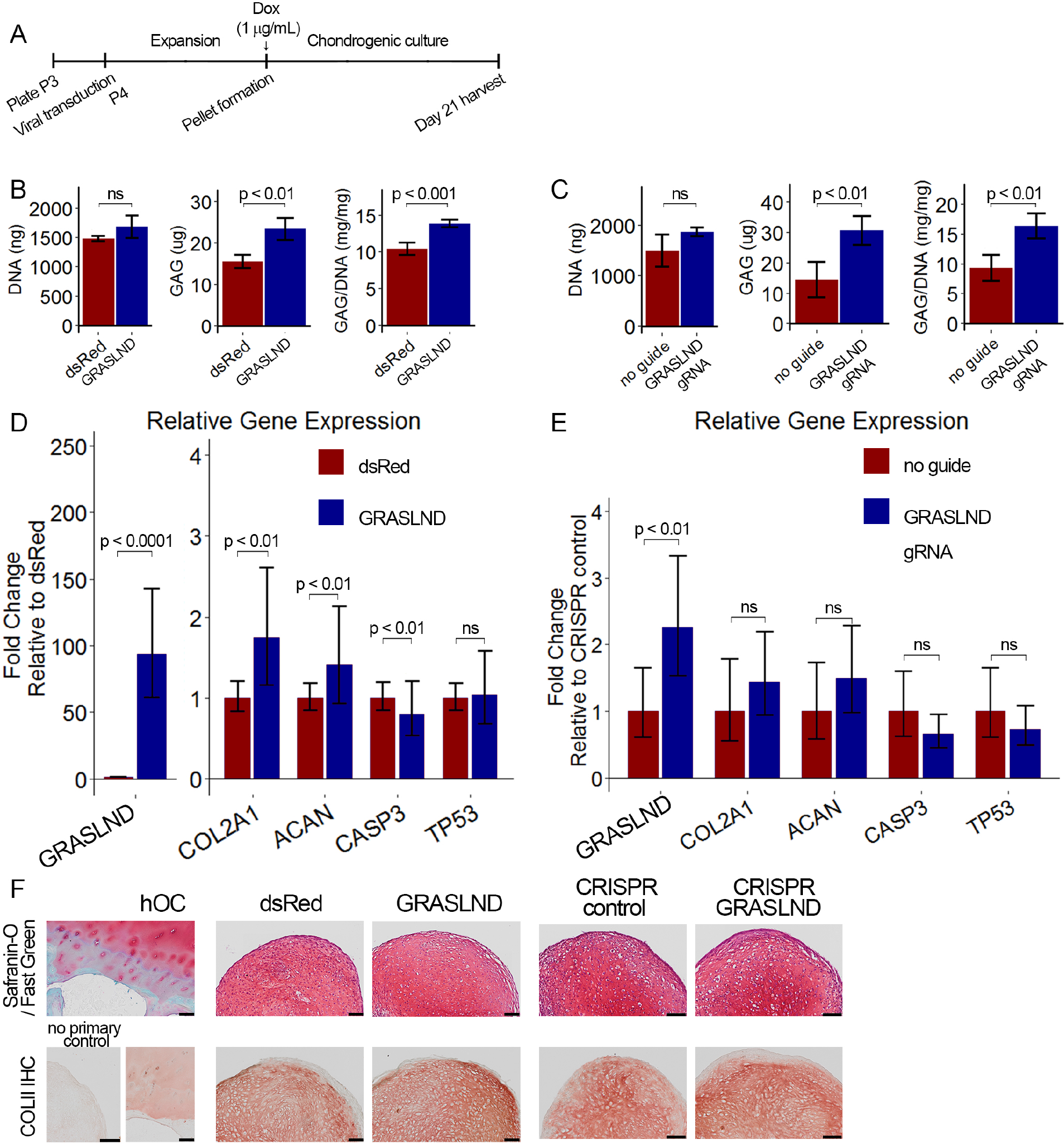
*GRASLND* enhances chondrogenesis. (A) Experimental timeline. (B,C) Biochemical analyses of day 21 MSC pellets (n=4). Welch’s t-test. (D,E) qRT-PCR analyses of day 21 MSC pellets (n=5 in D; n=6 in E). Welch’s t-test. (F) Representative histological images of day 21 MSC pellets. *COLII IHC: collagen type II immunohistochemistry. hOC: human osteochondral control.* Scale bar = 100 µm. (B,D,F) Transgene ectopic expression of *GRASLND*. (C,E,F) CRISPR-dCas9-VP64 induced activation of *GRASLND*. ns: not significant (p > 0.05).

To determine whether *GRASLND* would improve chondrogenesis at lower doses of growth factor or at earlier time points, we compared DNA and GAG levels from pellets cultured under different TGF-β3 concentrations on day 7, day 14, and day 21 (Figure S4 D-F). In agreement with our knockdown data, DNA content was unaffected. On the other hand, increases in GAG were observed at higher doses and at later time points, especially at 10 ng/mL of TGF-β3. It appears that an elevated level of *GRASLND* alone was not sufficient to enhance GAG deposition, and *GRASLND* may act in concert with other downstream effectors, which were not present at lower doses of TGF-β3 or at earlier time points in the process.

Elevated levels of *GRASLND* resulted in higher amounts of GAG deposition (p < 0.001) (Figure 4B), consistent with our data on the gene expression level (Figure 4D). We observed a slight increase in chondrogenic markers (*COL2A1*, *ACAN*), and a slight decrease in the apoptotic marker *CASP3*, while cellular senescence was not different between the two groups (*TP53*) (Figure 4D). Histologically, pellets derived from dsRed-transduced MSCs exhibited normal GAG and collagen type II staining, indicating successful chondrogenesis. The control pellets were indistinguishable from those derived from *GRASLND*-transduced MSCs (Figure 4F), albeit macroscopically smaller at the time of harvest.

These findings were further confirmed using CRISPR-dCas9-VP64 mediated activation of endogenous *GRASLND*. This system had been previously utilized to upregulate various transcription factors that efficiently induce embryonic fibroblasts into neurons (43, 44). After screening eleven synthetic gRNAs, we selected the one with highest activation level (Figure S5). When *GRASLND* was transcriptionally activated with CRISPR-dCas9, chondrogenesis was enhanced as evidenced by elevated amount of GAG deposition (p < 0.01); DNA amount may also be slightly increased, albeit not statistically significant (Figure 4C). Similar trends were detected by qRT-PCR (Figure 4E) and histology (Figure 4F). It is worth noting that CRISPR-dCas9 mediated activation only resulted in a moderate up-regulation of *GRASLND* relative to transgene ectopic expression (2-fold vs 100-fold). However, the functional outcome was more pronounced with CRISPR-dCas9. We observed approximately 50% increase in the level of GAG produced when normalized to DNA (9.4 ± 2.19 mg/mg vs 16.3 ± 2.08 mg/mg), compared to 30% detected with ectopic expression (10.5 ± 0.84 mg/mg vs 13.9 ± 0.52 mg/mg).

### *GRASLND* inhibits type II interferon signaling potentially by binding to EIF2AK2 and protects engineered cartilage from interferon

To decipher the potential signaling pathways involved, we chondrogenically induced MSCs in the presence or absence of *GRASLND,* and then utilized RNA-seq to compare the global transcriptomic changes between two conditions. As expected, *GRASLND* depletion resulted in impaired expression of chondrocyte-associated genes such as *TRPV4* and *COL9A2* (top 20 downregulated genes ranked by adjusted p-values) (Figure 5A). Skeletal system development and extracellular matrix organization were among the pathways most affected by the knockdown (Figure 5B). Surprisingly, pathways pertaining to interferon response were highly enriched in the upregulated gene list upon silencing of *GRASLND*. The top 20 upregulated genes involved many IFN downstream targets (*MX2, IFI44, IFI44L, IFITM1, IFI6, IFIT1, STAT1, MX1, IFIT3, OAS3, OAS2*), with both type I (IFN-α, IFN-β) and type II (IFN-γ) found to be enriched in our gene ontology analysis (Figure 5B). Furthermore, upregulated genes were also found to exhibit DNA binding motifs for transcription factors of the IFN pathways: STAT1, STAT2, IRF1, IRF2 (Table 2). A full list of differentially expressed genes is provided in Supplementary Materials. Further bioinformatic analyses created a network of potential transcription regulators as well as gene ontology terms for the upregulated gene cohort as a result of *GRASLND* silencing (Figure 5C). Taken together, *GRASLND* may potentially act to suppress the activities of these transcription factors, as a result affecting IFN signaling pathways during chondrogenesis.

**Figure 5:**
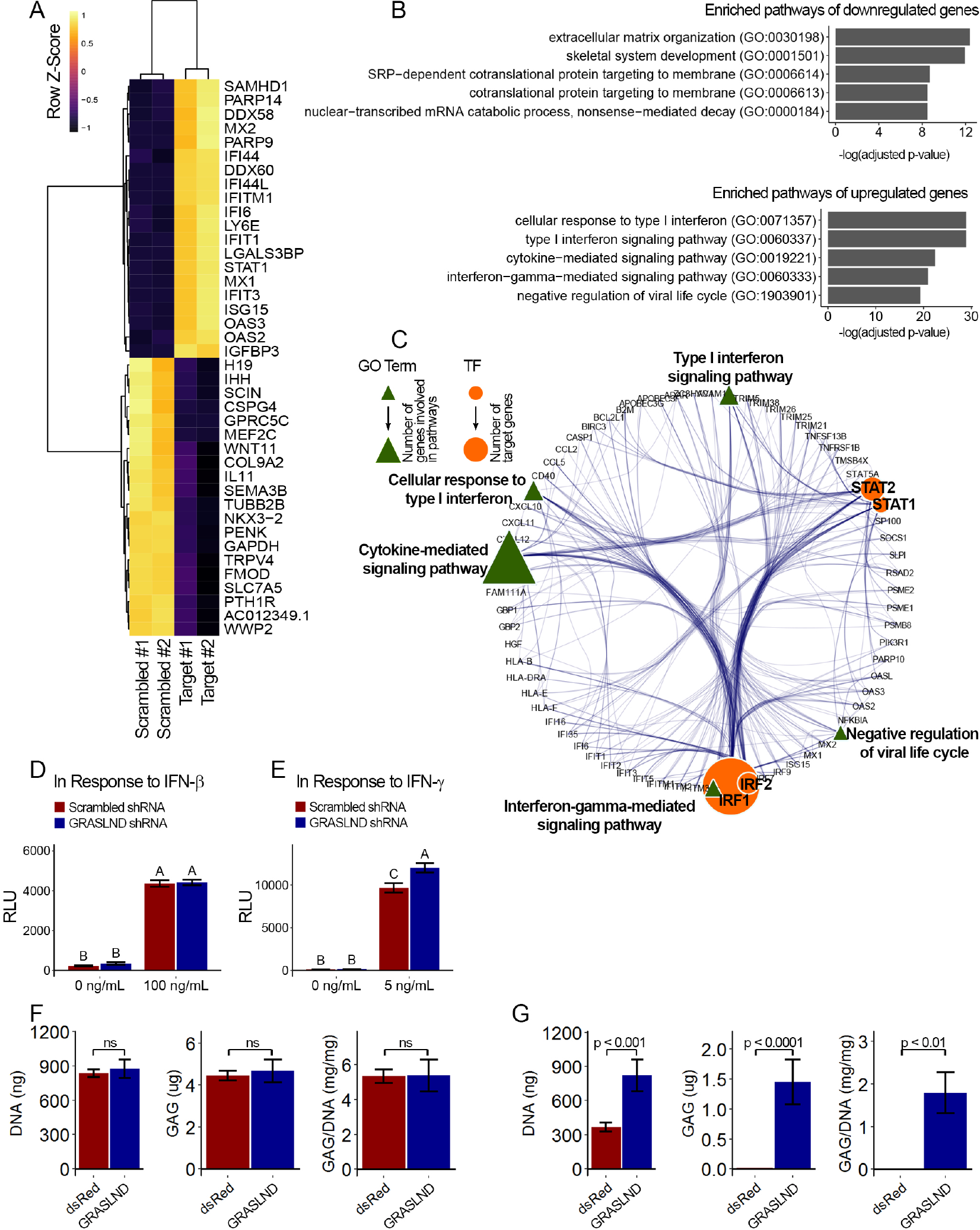
*GRASLND* suppresses interferon type II signaling. (A) Top 20 up- and down-regulated genes in *GRASLND* KD pellets compared to scrambled controls. (B) Gene ontology analysis of affected pathways. (C) Upregulated targets and related gene ontology terms and potential transcription factors. (D,E) Luciferase reporter assays on MSCs transduced with: (D) ISRE promoter element (n=3), or (E) GAS promoter element (n=3). Two-way ANOVA followed by Tukey post-hoc test (α=0.05). Groups of different letters are statistically different. (F) Biochemical assays on MSC-derived pellets cultured under 100 ng/mL of IFN-β (n=4). (G) Biochemical assays on MSC-derived pellets cultured under 5 ng/mL of IFN-γ (n=6). Welch’s t-test. *ns: not significant*.

**Table 2:**
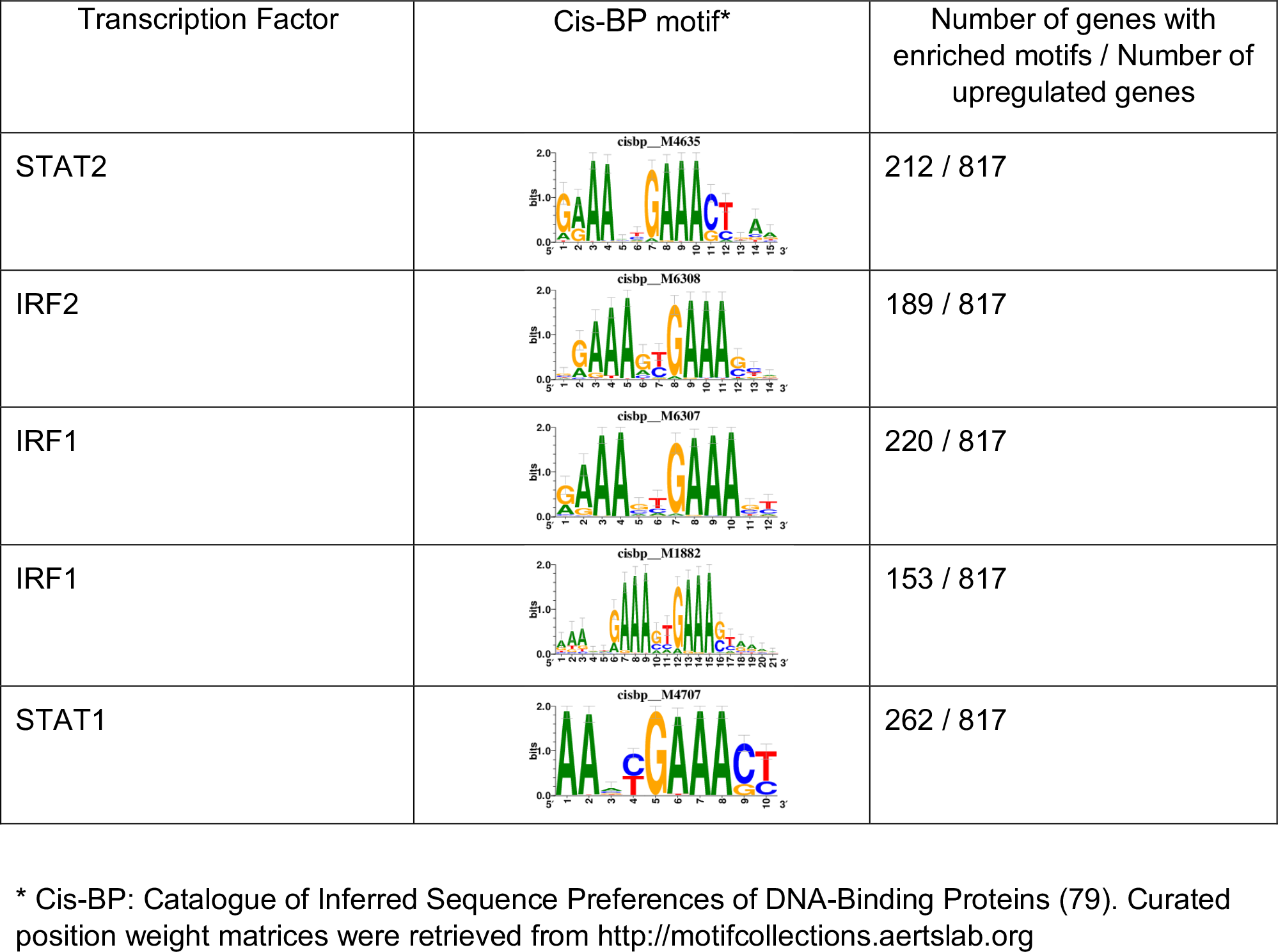
*Top 5 enriched Cis-BP motifs and associated transcription factors for upregulated genes upon GRASLND knockdown*

| Transcription Factor | Cis-BP motif* | Number of genes with enriched motifs / Number of upregulated genes |
| --- | --- | --- |
| STAT2 |  | 212 / 817 |
| IRF2 |  | 189 / 817 |
| IRF1 |  | 220 / 817 |
| IRF1 |  | 153 / 817 |
| STAT1 |  | 262 / 817 |

\* Cis-BP: Catalogue of Inferred Sequence Preferences of DNA-Binding Proteins (79). Curated position weight matrices were retrieved from http://motifcollections.aertslab.org

To further confirm this relationship, we performed luciferase reporter assays for interferon signaling upon *GRASLND* knockdown. Utilizing specific reporter constructs, we were able to determine whether *GRASLND* acted on type I or type II IFN. Our results indicated that decreased level of *GRASLND* led to heightened type II (IFN-γ) (Figure 5E) response but not type I (IFN-β) (Figure 5D). Importantly, luminescence activities between scrambled control and *GRASLND* knockdown were indistinguishable from each other in basal, IFN-free conditions. This indicates that at basal level, the two groups responded similarly to lentiviral transduction, and the observed difference in IFN signal was a consequence of *GRASLND* downregulation.

Since *GRASLND* was expressed in the cytoplasm (Figure 2C), we hypothesized that it is part of an RNA-protein complex. To test this, we performed an RNA pull down assay, followed by mass spectrometry. Here, streptavidin beads were used as control, or conjugated to sense or antisense strands of *GRASLND*. Naked or conjugated beads were then incubated with lysates from day 21 pellets, from which bound proteins were eluted for further analyses. We found that Interferon-Induced Double-Stranded RNA-Activated Protein Kinase (EIF2AK2) peptides were detected at elevated levels in sense samples as compared to antisense controls (p < 0.05); peptides were undetected in naked bead controls. Subsequent RNA pull-down followed by western blot confirmed EIF2AK2 as a binding partner of *GRASLND* (Figure S6). We detected an increased level of EIF2AK2 bound to the sense strand of *GRASLND* relative to the antisense or the pellet lysate control. We speculate that this association of *GRASLND* RNA to EIF2AK2 could potentially result in downregulation of IFN-γ signaling.

Interestingly, by mining a published microarray database (GSE57218) (45), we found that IFN-related genes were highly elevated in cartilage tissues of osteoarthritis patients: *STAT1, IFNGR2, NCAM1, MID1* (Figure S7A). Since the microarray did not contain probes for *GRASLND*, no information on its expression could be extracted. In addition, we identified another independent study that reported changes in the transcriptomes of intact and damaged cartilage tissues (E-MTAB-4304) (46). Similarly, a cohort of IFN-related genes were also upregulated in damaged cartilage, especially *STAT1* and *IFNGR1* (Figure S7B). Interestingly, we identified a negative correlation between *GRASLND* and a few IFN related genes (*IFNGR1*, *ICAM1*) in damaged cartilage (Figure S7C). Therefore, we proposed that *GRASLND* may possess some therapeutic potential through suppressing IFN signaling in osteoarthritis. To evaluate this possibility, we implemented the use of the *GRASLND* transgene in engineered cartilage cultured under IFN addition (100 ng/mL of IFN-β or 5 ng/mL of IFN-γ). We determined doses of IFN-*β* and IFN-γ by selecting the lowest concentration at which day 21 pellets exhibited GAG loss compared to no IFN control. Consistent with luciferase reporter assays, the protective effect of *GRASLND* was observed upon IFN-γ challenge but not IFN-β (Figure 5F, G). However, we observed a reduced level of GAG production compared to normal conditions, suggesting that *GRASLND* can protect the ECM from degradation, but not completely to control levels.

### *GRASLND* enhanced the chondrogenesis of adipose-derived stem cells

To determine if the function of *GRASLND* is unique to MSCs or present in other adult stem cells, we addressed whether modulating *GRASLND* expression could also improve chondrogenesis of adipose stem cells (ASCs). We observed an increase in GAG production when *GRASLND* was overexpressed in ASCs compared to control (p < 0.0001) (Figure 6A), although *ACAN* levels were not significantly increased. Importantly, *COL2A1* expression was significantly elevated (∼ 5-fold) with overexpression of *GRASLND* (Figure 6B). Histologic examination of the engineered cartilage showed a similar level of collagen type II in pellets with *GRASLND* overexpression compared to the dsRed control (Figure 6C). Based on these data, it appears that *GRASLND* utilized the same mechanism across these two cell types, asserting a pan effect on potentiating their chondrogenic capabilities.

**Figure 6:**
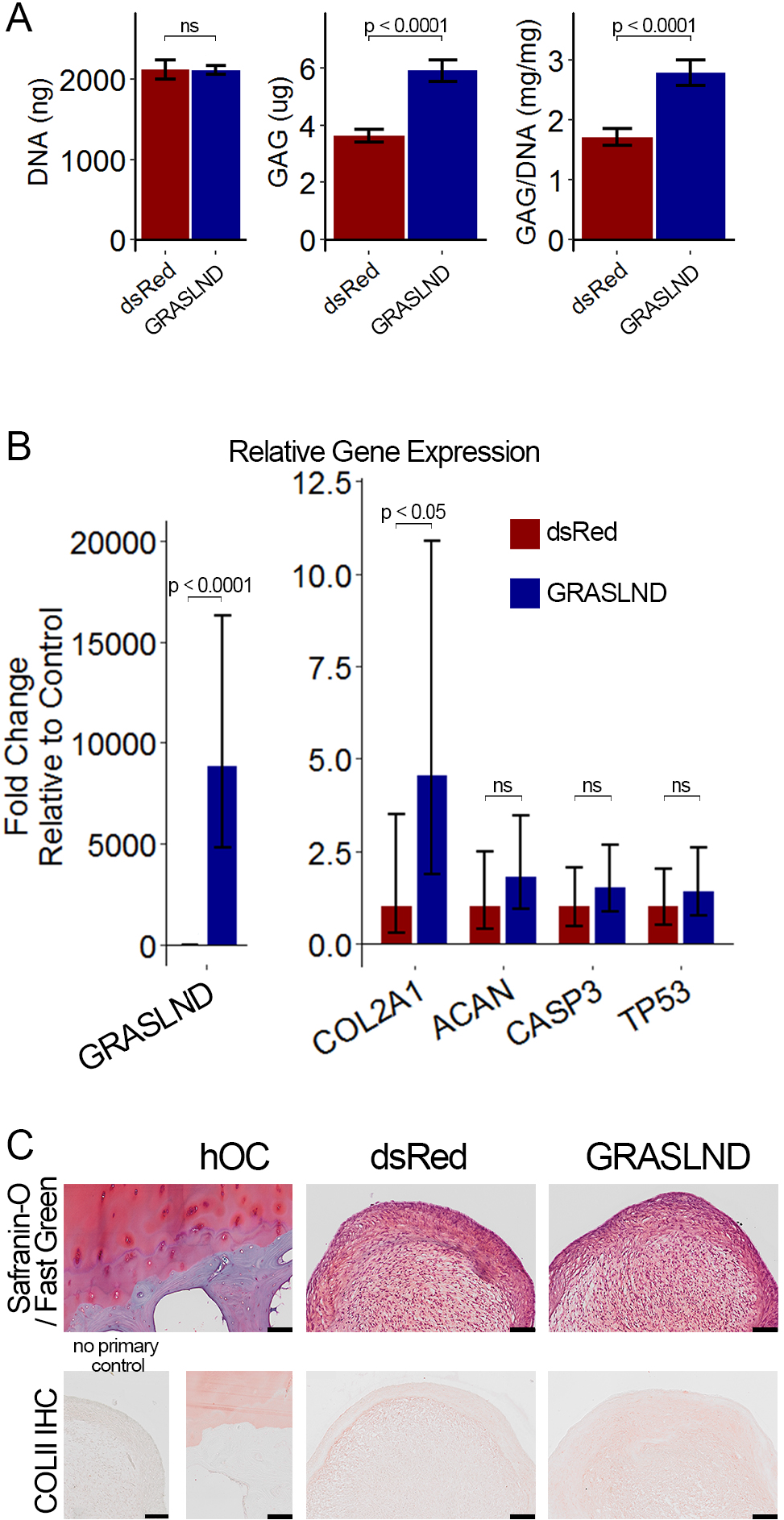
*GRASLND* enhances chondrogenesis in adipose-derived stem cells. (A) Biochemical analyses (n= 5). (B) qRT-PCR analyses (n=6). (C) Representative histological images of day 21 ASC pellets. *COLII IHC: Collagen type II immunohistochemistry. hOC: Human osteochondral control.* Scale bar = 100 µm. Welch’s t-test. ns: not significant.

## Discussion

Here, we identified and demonstrated the first functional study of lncRNA *GRASLND,* which acts to enhance stem cell chondrogenesis. Knockdown of *GRASLND* via shRNA inhibited chondrogenesis, whereas ectopic transgene or CRISPR-based overexpression of *GRASLND* enhanced chondrogenesis of MSCs and ASCs. Pathway analysis revealed a link between *GRASLND* and IFN-γ signaling pathway in this process, which was confirmed by the identification of EIF2AK2 as its binding partner. Unfortunately, lack of a known murine homolog makes it difficult to study *GRASLND in vivo,* and thus future studies may require *GRASLND* transgenic models in primate species.

In the context of the musculoskeletal system, IFN is mostly recognized for its role in bone development and homeostasis (23–27, 30), myogenesis (29, 47, 48), as well as its crosstalk with TGF-β in wound healing (49). Notably, IFN-γ has been suggested to inhibit collagen synthesis in dermal fibroblasts, myofibroblasts, and articular chondrocytes (49–53). Furthermore, the JAK/STAT pathway, which involves IFN downstream effectors, has also been shown to inhibit chondrocyte proliferation and differentiation (28, 31). Here, we found that *GRASLND* acts to suppress the IFN mechanism. In addition, we also present evidence indicating an interaction between *GRASLND* and EIF2AK2 (also referred to as PKR). Canonically a crucial player in protein synthesis, PKR has also been reported to control STAT signaling by directly binding to and preventing its association with DNA for gene activation (54, 55). Additionally, several studies have suggested that highly structured, single stranded RNA can also activate PKR via its double stranded RNA binding domains (dsDRBs) (56–60). Our RNA-seq data suggested that upon *GRASLND* knockdown, a cohort of downstream targets of STATs were upregulated.

Based on the presence of DNA binding motifs in investigated targets, we identified both STAT1 and STAT2 as potential regulators of genes disrupted by GRASLND knockdown. However, our luciferase reporter assays pointed towards a mechanism in IFN type II (gene activation by STAT1 homodimer) rather than type I (gene activation by STAT1/STAT2 heterodimer). Thus, we hypothesized that *GRASLND* could form a secondary structure to bind and activate PKR, which in turn inhibits STAT1-related transcriptional function. This mechanism supports the hypothesis that modulation of IFN-γ via the JAK/STAT pathway, achieved by the *GRASLND/*PKR RNA-protein complex, is important for cellular proliferation and differentiation during chondrogenesis.

Upregulation of IFN has also been implicated in arthritis by several studies (61–64). Publicly available databases provide evidence corroborating similar patterns of IFN in degenerated cartilage. As *GRASLND* inhibits IFN, utilization of this lncRNA offers potential in both MSC cartilage tissue engineering and in OA treatment. As a proof of concept, we showed that *GRASLND* could enhance matrix deposition across cell types of origin, with and without interferon challenge *in vitro*. It would be interesting to next investigate whether *GRASLND* can protect cartilage from degradation in a milieu of pro-inflammatory cytokines *in vivo*.

Since lentivirus was employed to manipulate the expression of *GRASLND*, it is possible our observations were confounded by the cellular response to viral infection. However, our luciferase reporter assays demonstrated that basal luminescence levels (with no interferon supplementation) between the scrambled controls and the shRNA treatments were indistinguishable. This suggests that altered levels of interferon signaling can be attributed to experimentally varied levels of *GRASLND* and not to the presence of lentivirus. Our data indicate that *GRASLND* acts through type II rather than type I IFN. We found that 5 ng/mL of IFN-γ was still more detrimental to chondrogenic constructs compared to 100 ng/mL of IFN-β. One potential explanation for this phenomenon may be the skewed distribution of available surface receptors between type I and type II (IFNAR vs IFNGR). Indeed, MSCs express a much lower level of *IFNAR2* compared to *IFNAR1*, *IFNGR1*, or *IFNGR2* (both in GSE109503 (36) and in GSE129985 (this manuscript)). As these receptors function as heterodimers (32, 34), response to type I may be stunted due to IFNAR2 deficiency.

Furthermore, we showed that a modified CRISPR-dCas9 system could successfully be used for endogenous transcriptional activation of lncRNA. This system had been previously used in other cell types to regulate expression of both protein-coding and non-coding genes (43, 44, 65, 66). We showed that CRISPR may be more effective than transgene expression, as indicated by a larger increase in GAG production, despite lower levels of overall gene activation. As *GRASLND* does not regulate *RNF144A,* it is evident that *GRASLND* acts *in trans*. However, we speculate the CRISPR-dCas9 system could also be useful for gain of function studies to investigate lncRNAs acting *in cis*, as well as lncRNAs that are difficult to obtain via molecular cloning due to their secondary structures, high repeated sequence or GC-rich content.

In conclusion, we have identified *GRASLND* as an important regulator of chondrogenesis. *GRASLND* acts downstream of SOX9 and enhances cartilage-like matrix deposition in stem cell-derived constructs. Moreover, *GRASLND* functions to suppress IFN via PKR, and as a result induces adult stem cells towards a more chondrocyte-like lineage. It is likely that the *GRASLND*/PKR RNA-protein complex may inhibit STAT1 transcriptional activity. We propose that *GRASLND* can potentially be applied therapeutically for both cartilage tissue engineering and for the treatment of OA.

## Materials and Methods

### Cell culture

Bone marrow was obtained from discarded and de-identified waste tissue from adult bone marrow transplant donors in accordance with the Institutional Review Board of Duke University Medical Center. Adherent cells were expanded and maintained in expansion medium: DMEM-low glucose (Gibco), 1% Penicillin/streptomycin (Gibco), 10% fetal bovine serum (FBS) (ThermoFisher), and 1 ng/mL basic fibroblast growth factor (Roche) (67).

Adipose derived stem cells (ASCs) were purchased from ATCC (SCRC-4000) and cultured in complete growth medium: Mesenchymal stem cells basal medium (ATCC PCS-500-030), mesenchymal stem cell growth kit (ATCC PCS-500-040) (2% FBS, 5 ng/mL basic recombinant human FGF, 5 ng/mL acidic recombinant human FGF, 5 ng/mL recombinant human EGF, 2.4 nM L-alanyl-L-glutamine), 0.2 mg/mL G418.

### Plasmid construction

#### shRNA

Short hairpin RNA (shRNA) sequences for specific genes of interest were designed with the Broad Institute GPP Web Portal (68). For each gene, six different sequences were selected for screening, after which the two most effective were chosen for downstream experiments in chondrogenic assays. Selected shRNAs were cloned into a modified lentiviral vector (Addgene #12247) using MluI and ClaI restriction sites, as described previously (69). A complete list of effective shRNA sequences is presented in Table S1.

### Transgene overexpression of *GRASLND*

A derivative vector from modified TMPrtTA (3, 70) was created with NEBuilder® HiFi DNA Assembly Master Mix (New England Biolabs). Backbone was digested with EcoRV-HF (New England Biolabs) and PspXI (New England Biolabs). The following resultant fragments were amplified by polymerase chain reaction and assembled into the digested plasmid: Tetracycline responsive element and minimal CMV promoter (TRE/CMV), Firefly luciferase, bGH poly(A) termination signal (BGHpA). Primers and plasmids for cloning are provided in Table S2.

The full sequence of *GRASLND* transcript variant 1 (RefSeq NR_033997.1) was synthesized by Integrated DNA Technologies, Inc. *GRASLND* or the Discosoma sp. red fluorescent protein coding sequence (dsRed) were cloned into the above derivative tetracycline inducible plasmid with NEBuilder® HiFi DNA Assembly Master Mix (New England Biolabs) at NheI and MluI restriction sites (pLVD-*GRASLND* and pLVD-dsRed). Amplifying primers are provided in Table S2.

### CRISPR-dCas9 activation of *GRASLND*

Guide RNA sequences were designed using the UCSC genome browser (http://genome.ucsc.edu/) (71), integrated with the MIT specificity score calculated by CRISPOR and the Doench efficiency score (72, 73). Oligonucleotides (IDT, Inc) were phosphorylated, annealed, and ligated into the pLV-hUbC-dCas9-VP64 lentiviral transfer vector (Addgene #53192) previously digested at BsmBI restriction sites (74). Eleven potential guide RNA sequences were selected and screened for their efficacy, and the gRNA with the highest activation potential was chosen for further experiments (Figure S5). The synthetic gRNA used in all CRISPR-dCas9 activation experiments has the following sequence: 5′-CCACTGGGGATAGTTCCCTG-3′.

### Chondrogenesis assay

MSCs or ASCs were digested in 0.05% Trypsin-EDTA (Gibco), and trypsin was inactivated with 1.5X volume of expansion medium. Dissociated cells were centrifuged at 200 × g for 5 minutes, and supernatant was aspirated. Subsequently, cells were washed in pre-warmed DMEM-high glucose (Gibco) three times, and resuspended at 5 × 10^5^ cells/mL in complete chondrogenic medium: DMEM-high glucose (Gibco), 1% Penicillin/ streptomycin (Gibco), 1% ITS+ (Corning), 100 nM Dexamethasone (Sigma-Aldrich), 50 µg/mL ascorbic acid (Sigma-Aldrich), 40 µg/mL L-proline (Sigma-Aldrich), 10 ng/mL rhTGF-β3 (R&D Systems). Five hundred µL of the above cell mixture was dispensed into 15 mL conical tubes, and centrifuged at 200 × g for 5 minutes. Pellets were cultured at 37°C, 5% CO_2_ for 21 days with medium exchange every three days.

### Osteogenesis and adipogenesis assays

MSCs were plated at 2 × 10^4^ cells/well in 6-well plates (Corning) and cultured for 4 days in MSC expansion medium, followed by induction medium for 7 days. Osteogenic induction medium includes: DMEM-high glucose (Gibco), 10% FBS, 1% Penicillin/ streptomycin (Gibco), 10 nM Dexamethasone (Sigma-Aldrich), 50 µg/mL ascorbic acid (Sigma-Aldrich), 40 µg/mL L-proline (Sigma-Aldrich), 10 mM β-glycerol phosphate (Chem-Impex International), 100 ng/mL rh-BMP2 (ThermoFisher). Adipogenic induction medium includes: DMEM-high glucose (Gibco), 10% FBS (ThermoFisher), 1% Penicillin/ streptomycin (Gibco), 1% ITS+ (Corning), 100 nM Dexamethasone (Sigma-Aldrich), 450 µM 3-isobutyl-1-methylxanthine (Sigma-Aldrich), 200 µM indomethacin (Sigma-Aldrich).

### Biochemical assays

Harvested pellets were stored at −20°C until further processing. Collected samples were digested in 125 µg/mL papain at 60°C overnight. DMMB assay was performed as previously described to measure GAG production (76). PicoGreen assay (ThermoFisher) was performed to measure DNA content following manufacture’s protocol.

### Immunohistochemistry and histology

Harvested pellets were fixed in 4% paraformaldehyde for 48 hours, and processed for paraffin embedding. Samples were sectioned at 10 µm thickness, and subjected to either Safranin O – Fast Green standard staining (77) or to immunohistochemistry of collagen type II (Developmental Studies Hybridoma Bank, University of Iowa; #II-II6B3). Human osteochondral sections were stained simultaneously to serve as positive control. Sections with no primary antibodies were used as negative control for immunohistochemistry.

### RNA fluorescence in situ hybridization (RNA FISH)

Harvested pellets were snap frozen in Tissue-Plus O.C.T. Compound (Fisher HealthCare) and stored at −80°C until further processing. Samples were sectioned at 5 µm thickness and slides were stored at −80°C until staining. Probe sets for RNA FISH were conjugated with Quasar® 670 dye, and were synthesized by LGC Biosearch Technologies and listed in Table S3 (*RNF144A-AS1*). GAPDH probe set was pre-designed by the manufacturer. Staining was carried out according to manufacturer’s protocol for frozen tissues. Slides were mounted with Prolong Gold anti-fade mountant with DAPI (ThermoFisher) and imaged with the Virtual Slide Microscope VS120 (Olympus) at lower magnification and with the confocal microscope (Zeiss) at higher magnification.

### RNA isolation and quantitative RT-PCR

Norgen Total RNA Isolation Plus Micro Kits (Norgen Biotek) were used to extract RNA from pellet samples and Norgen Total RNA Isolation Plus Kits (Norgen Biotek) were used for all other RNA isolation. For monolayer, cells were lysed in buffer RL and stored at −20°C until further processing. For pellets, harvested samples were snap frozen in liquid nitrogen and stored at −80°C until further processing. On day of RNA isolation, pellets were homogenized in buffer RL using a bead beater (BioSpec Products) at 2,500 oscillations per minute for 20 seconds for a total of three times. Subsequent steps were performed following manufacturer’s protocol.

Nuclear and cytoplasmic fractions from day 21 MSC pellets were separated with the NE-PER Nuclear and Cytoplasmic Extraction Reagents (ThermoFisher) following manufacturer’s protocol. Resulting extracts were immediately subjected to RNA isolation using the Norgen Total RNA Isolation Plus Micro Kits (Norgen Biotek) by adding 2.5 parts of buffer RL to 1 part of extract. Subsequent steps were carried out following manufacturer’s protocol.

Reverse transcription by Superscript VILO cDNA master mix (Invitrogen) was performed immediately following RNA isolation. cDNA was stored at −20°C until further processing. qRT-PCR was carried out using Fast SyBR Green master mix (Applied Biosystems) following manufacturer’s protocol. A complete list of primer pairs (synthesized by Integrated DNA Technologies, Inc.) is reported in Table S4.

### Luminescence assay

MSCs were plated at 8.5 × 10^4^ cells per well in 24-well plates (Corning). Lentivirus carrying the response elements for type I (ISRE - #CLS-008L-1) or type II (GAS - #CLS-009L-1) upstream of firefly luciferase was purchased from Qiagen. Twenty-four hours post plating, cells were co-transduced with virus in the following groups: ISRE with scrambled shRNA, ISRE with *GRASLND* shRNA, GAS with scrambled shRNA, GAS with *GRASLND* shRNA. Twenty-four hours post-transduction, cells were rinsed once in PBS and fresh medium was exchanged.

Three days later, medium was switched to expansion medium with 100 ng/mL IFN-β (PeproTech) for wells with ISRE or with 5 ng/mL IFN-γ (PeproTech) for wells with GAS. MSCs were cultured for another 22 hours, and then harvested for luminescence assay using Bright-Glo Luciferase Assay System (Promega). Luminescence signals were measured using the Cytation 5 Plate reader (BioTek).

### Western blot

On day of harvest, cells were homogenized with complete lysis buffer in ice cold PBS: 10X RIPA buffer (Cell Signaling Technology), 100X phosphatase inhibitor cocktail A (Santa Cruz Biotechnology), 100X Halt^TM^ protease inhibitor cocktails (ThermoScientific). Lysates were subsequently centrifuged at 14,000 × g for 15 minutes at 4°C, and supernatants were collected and stored at −20°C until further processing. Western blot was serviced by RayBiotech with the following antibodies: primary anti-β-actin (RayBiotech), primary anti-RNF144A (Abcam), primary anti-PKR (RayBiotech) and secondary anti-rabbit-HRP (horse radish peroxidase) (RayBiotech).

### Statistical analyses

All statistical analyses were performed using R (78). Results from biochemical assays are depicted as mean ± SD. Results from qRT-PCR are depicted as fold-change with error bars calculated per Applied Biosystems manual instruction.

Additional methods are provided in supplemental information.

## Acknowledgments

We thank the Genome Technology Access Center at Washington University in St Louis, the Proteomics Core Laboratory, and the Hope Center Viral Vectors Core for their resources and support. The CRISPR-dCas9-VP64 system was a generous gift from Dr. Charles Gersbach. We also wish to thank Sara Oswald for providing assistance in technical writing of the manuscript. This work was supported by the Arthritis Foundation, NIH grants AR50245, AR48852, AG15768, AR48182, AR067467, AR057235, AR073752, the Nancy Taylor Foundation for Chronic Diseases, and the Collaborative Research Center of the AO Foundation, Davos, Switzerland.

## Author contributions

N.P.T.H, F.G designed research; N.P.T.H., C.C.G., J.L. and R.T. performed research and analyzed data; J.M.B., A.M., and B.Z. provided critical discussion and comments. N.P.T.H. and F.G. wrote the manuscript; all authors edited the manuscript.

**Accession number:** GSE129985

## Supplemental Information

### I. Supplemental Figures

**Figure S1:**
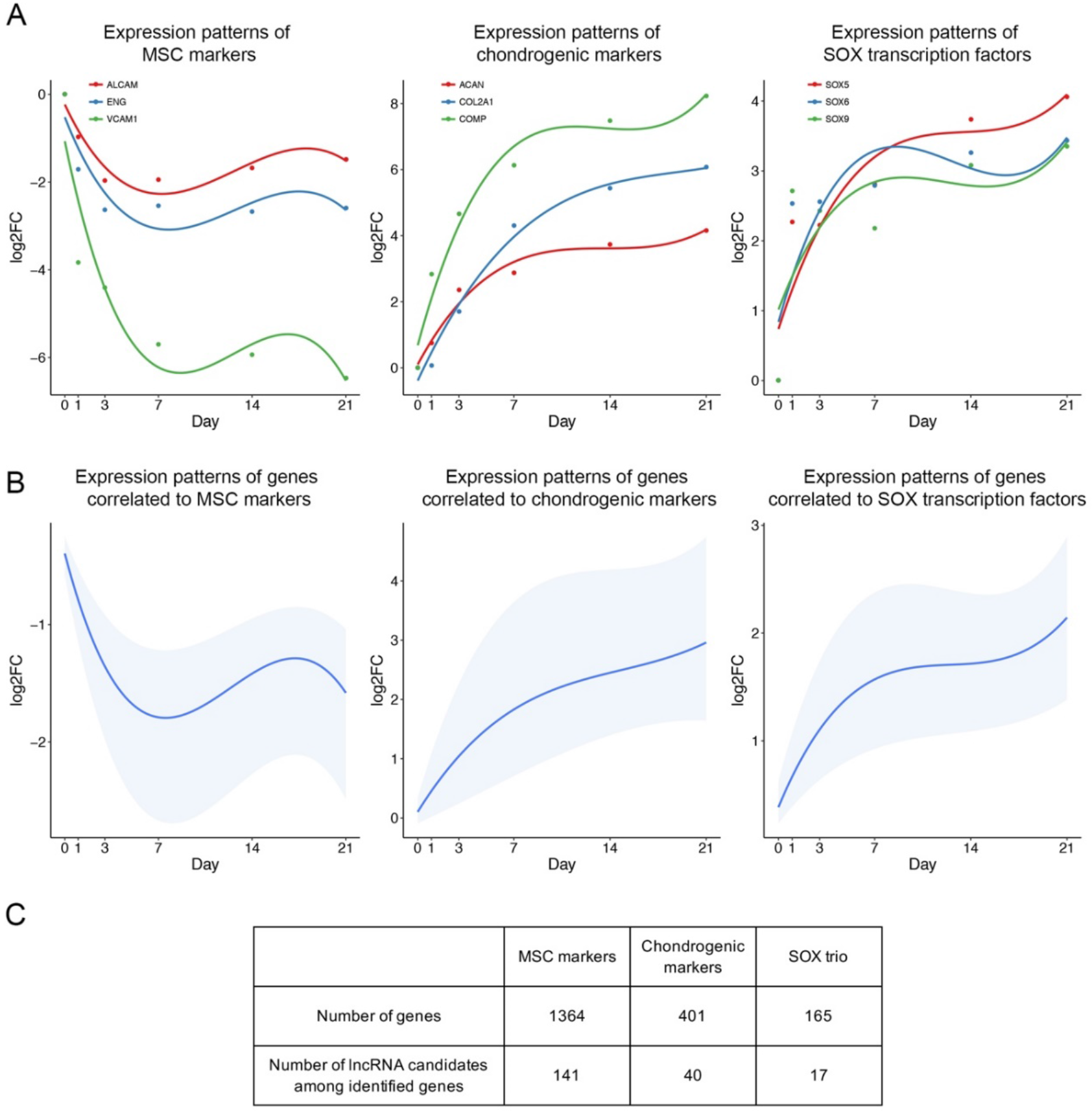
Identification of lncRNA candidates. LncRNAs whose expression patterns are correlated to crucial markers are of interest. (A) Expression patterns of previously identified MSC markers (left), chondrogenic markers (middle), and SOX transcription factors (right). Data retrieved from: GSE109503. (B) Expression patterns of correlated genes or lncRNAs (Pearson correlation > 0.9) to MSC markers (left), chondrogenic markers (middle), and SOX transcription factors (right). Lower boundary: 25^th^ percentile; upper boundary: 75^th^ percentile; blue line: median; of correlated gene set. (C) Number of correlated genes and correlated lncRNAs.

**Figure S2:**
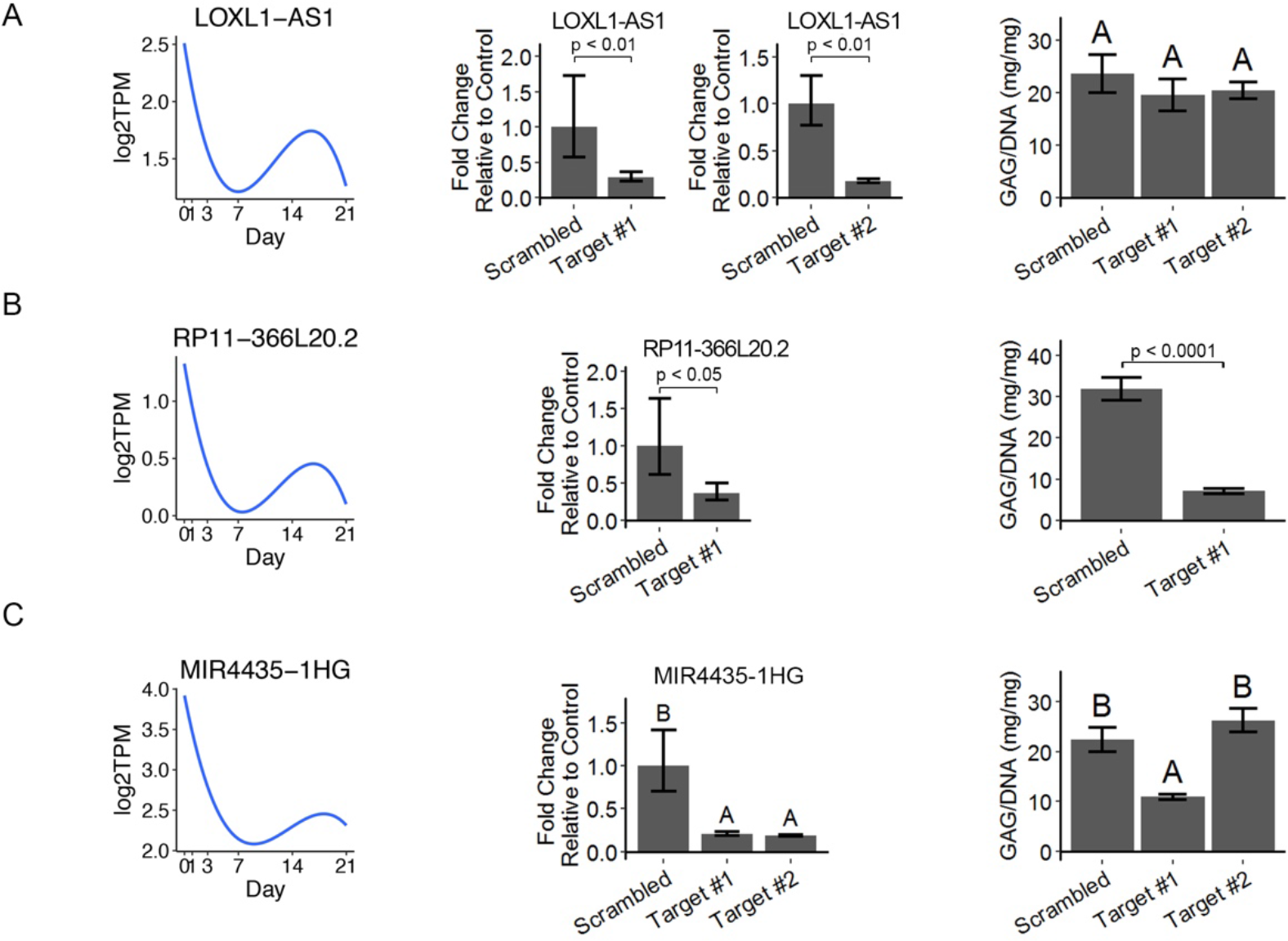
Functional validation of identified lncRNA candidates. **<u>Left</u>**: Expression pattern of candidates during chondrogenesis (GSE109503). Log2TPM: log-transformed value of transcripts per million (TPM). **<u>Middle</u>**: Efficiency of designed target shRNAs. Individual graphs indicate experiments on target number 1 and target number 2 were performed separately (n=3). Welch’s t-test on log transformed fold changes for A-B. One-way ANOVA with Tukey post-hoc test for C (α=0.05). Groups of different letters are statistically different. **<u>Right</u>**: Quantitative analysis of synthesized GAG matrix normalized to DNA amount. (A) LOXL1-AS1 (n=4-5). One-way ANOVA with Tukey post-hoc test. Groups of different letters are statistically different. (B) RP11-366L20.2 (n=3-5). Welch’s t-test. (C) MIR4435-1HG (n=3-4). One-way ANOVA with Tukey post-hoc test. Groups of different letters are statistically different.

**Figure S3:**
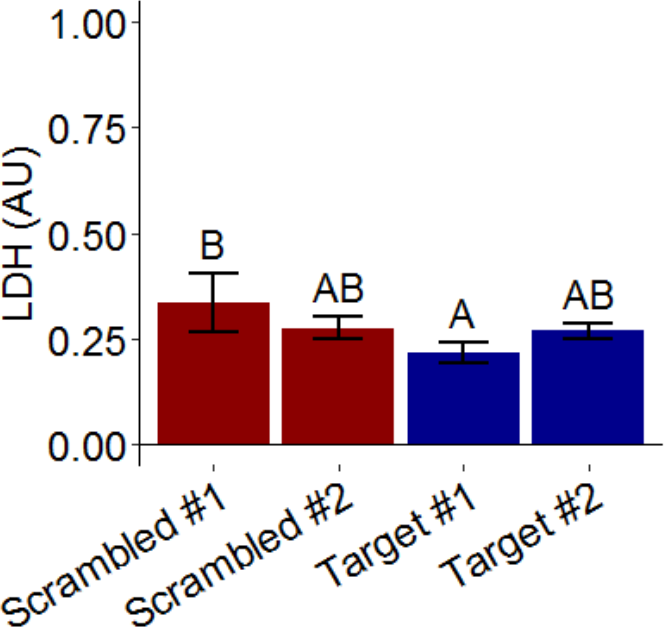
Cytotoxicity assay of target shRNAs. Cytotoxicity level was measured indirectly by the amount of released LDH (n = 4). One-way ANOVA followed by Tukey post-hoc test (α =0.05). Groups of different letters are statistically different from one another.

**Figure S4:**
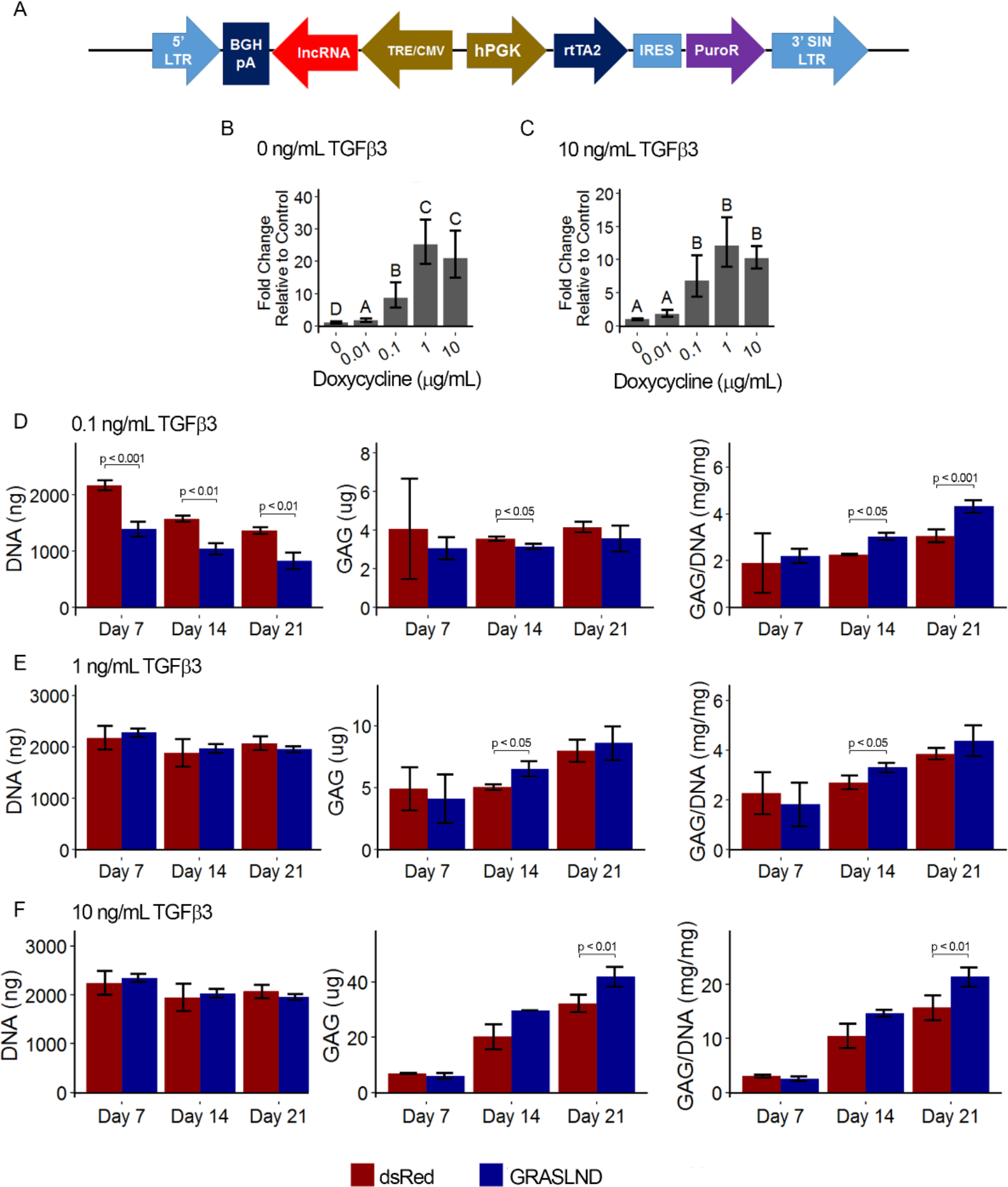
Effect of *GRASLND* overexpression across time points and tested doses. (A) Overview of designed lentiviral backbone. *GRASLND* is driven under a Doxycycline inducible promoter, and poly-adenylated with BGHpA signal. (B-C) Relative expression of *GRASLND* under different doses of Doxycycline (Dox). Dox is most potent at 1 μg/mL under both conditions of TGF-β3 (n=4). One-way ANOVA with Tukey post-hoc test (α=0.05). Groups of different letters are statistically different. (D-F) Biochemical analyses of MSC pellets cultured under chondrogenic condition with different doses of TGF-β3 (n = 3-4). Welch’s t-test. No bracket indicates the comparison is not significant. *LTR: Long terminal repeat; BGHpA: Bovine growth hormone polyadenylation signal; TRE/CMV: Tet responsible element fused with the minimal cytomegalovirus promoter; rtTA2: Reverse tetracycline-controlled transactivator 2; IRES: Internal ribosome entry site; PuroR: Puromycin N-acetyl-transferase; SIN: Self inactivating*.

**Figure S5:**
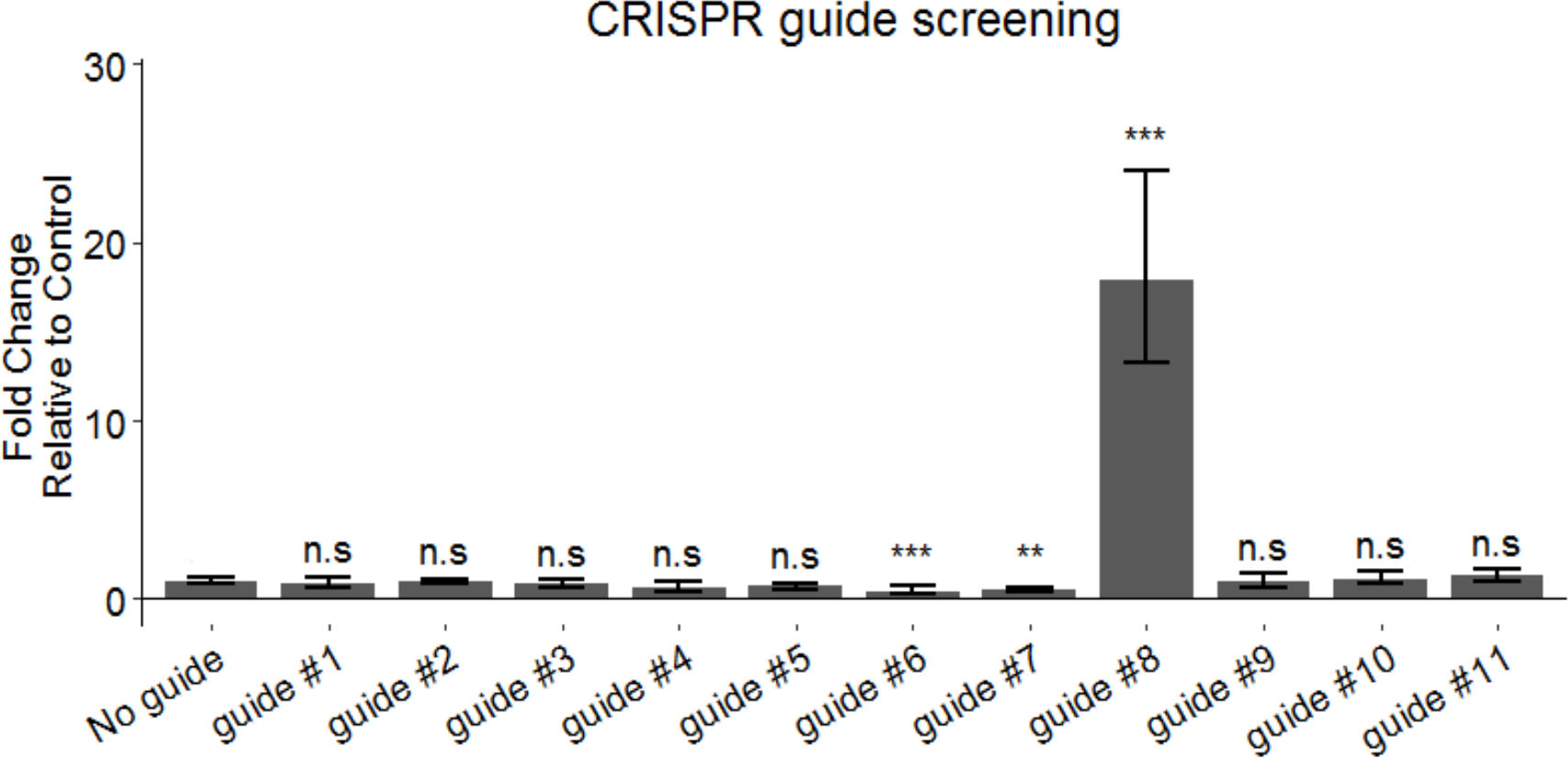
Synthetic guide RNA screening for efficient activation of endogenous *GRASLND*. Dunnett’s test compared to “No Guide” control. ns: not significant. ** p < 0.01; *** p < 0.001.

**Figure S6:**
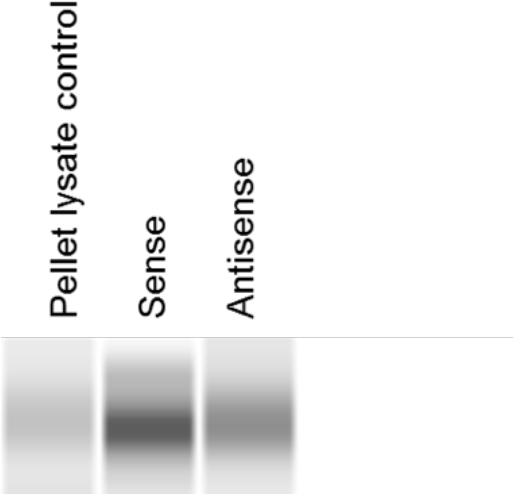
RNA pull-down followed by western blot confirmed EIF2AK2 as the binding partner of *GRASLND*.

**Figure S7:**
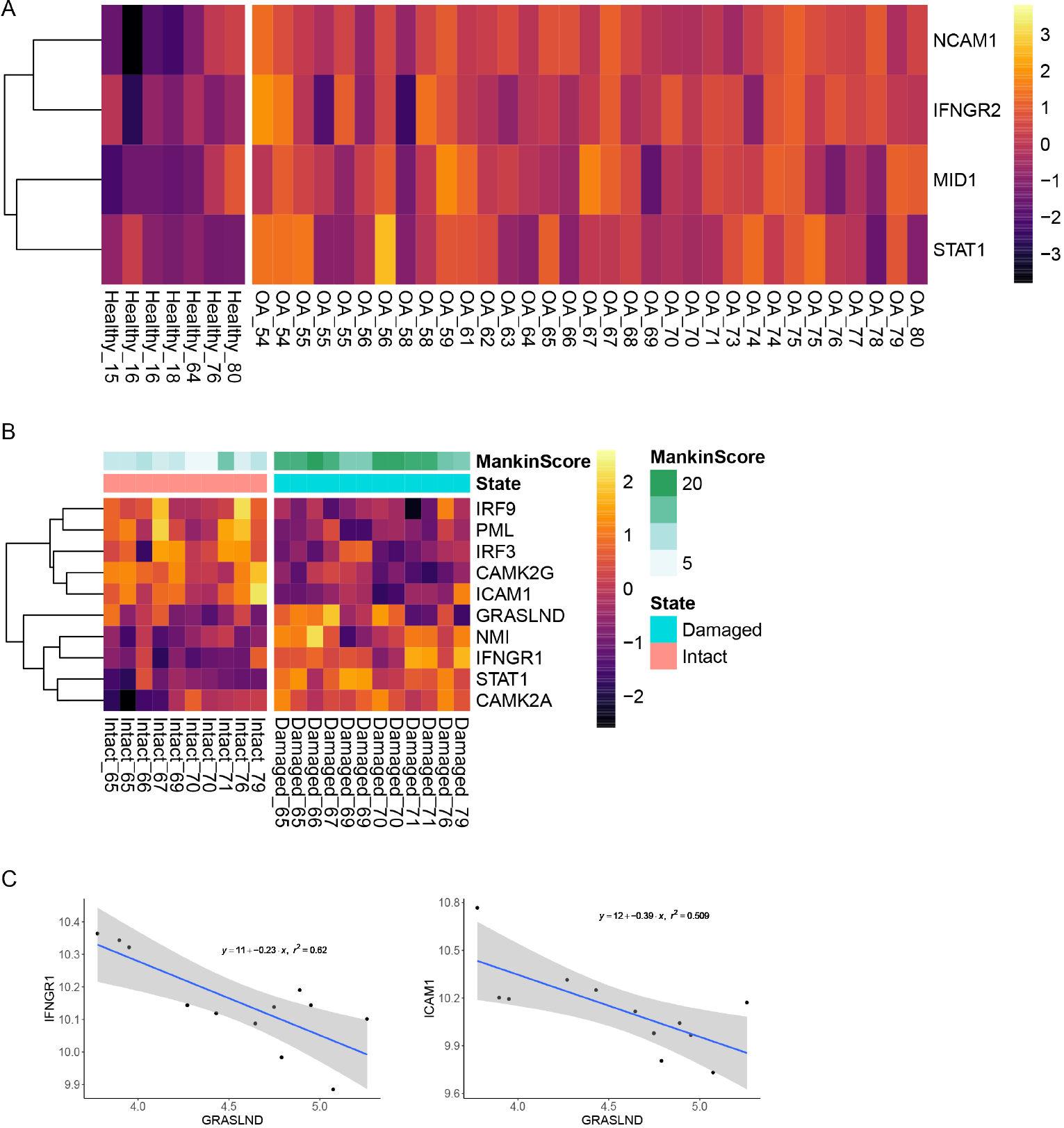
IFN signaling from other publicly available databases. (A) IFN signal was upregulated in OA patients. Samples are indicated as disease state followed by patients’ age. (A) IFN was upregulated while *GRASLND* was downregulated in damaged cartilage. Samples are indicated as cartilage site followed by patients’ age. (C) Inverse correlation between *GRASLND* and IFN-related genes.

### II. Supplemental Tables

**Table S1:**
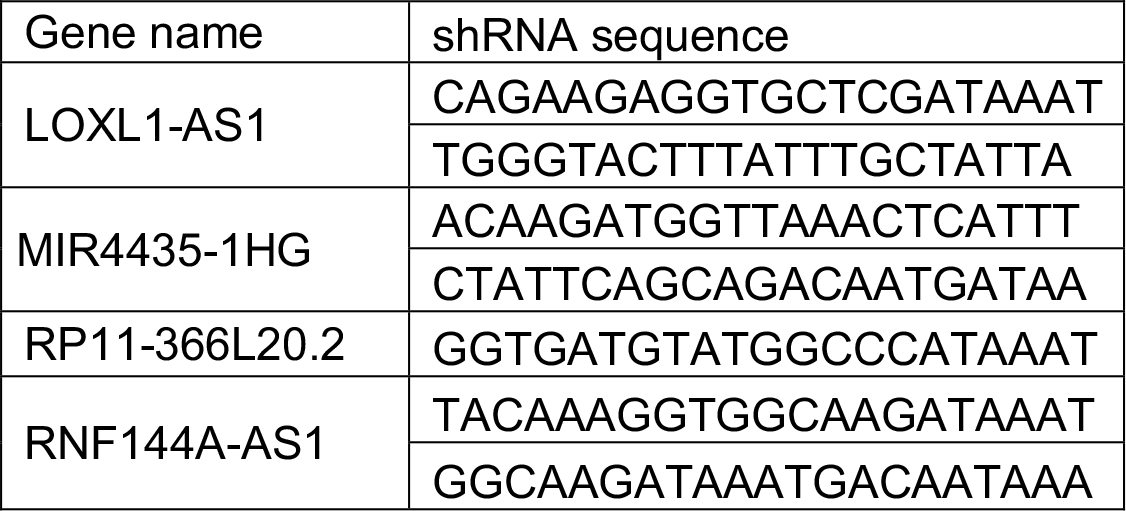
shRNA target sequences. 5′ – sequence – 3′

**Table S2:**
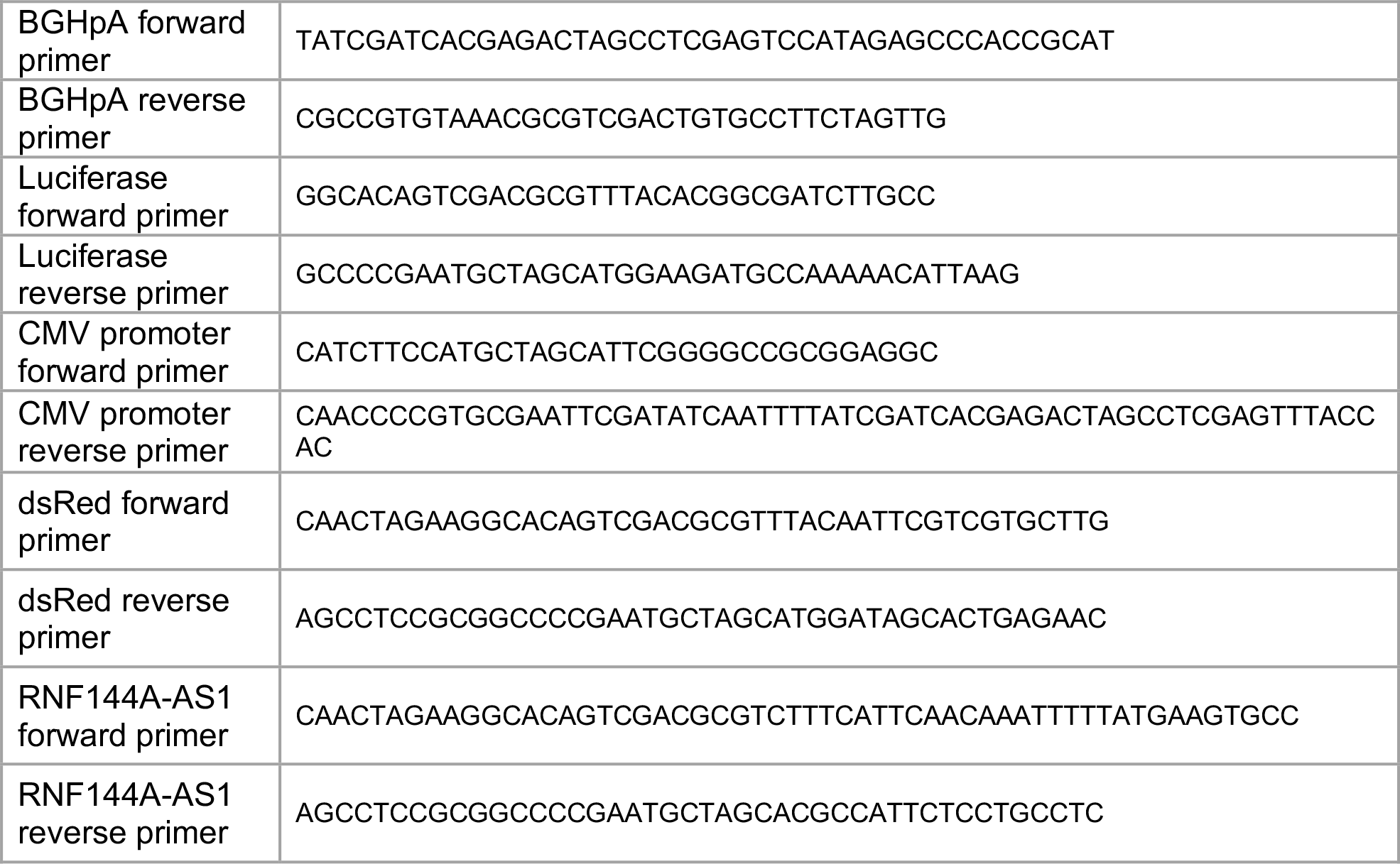
Cloning primer sequences. 5′ – sequence – 3′

**Table S3:**
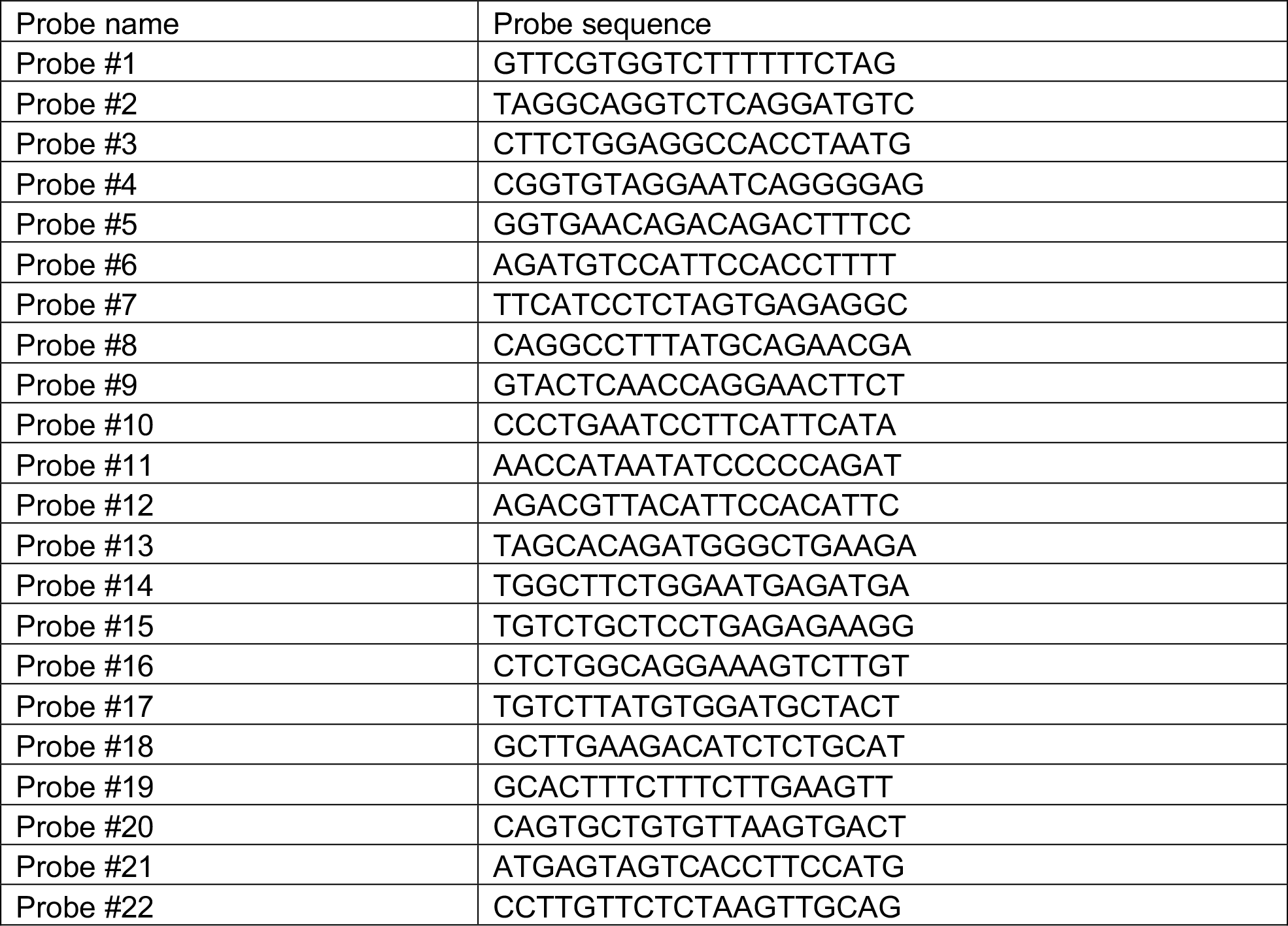
RNF144A-AS1 probe set sequences. 5′ – sequence – 3′

**Table S4:**
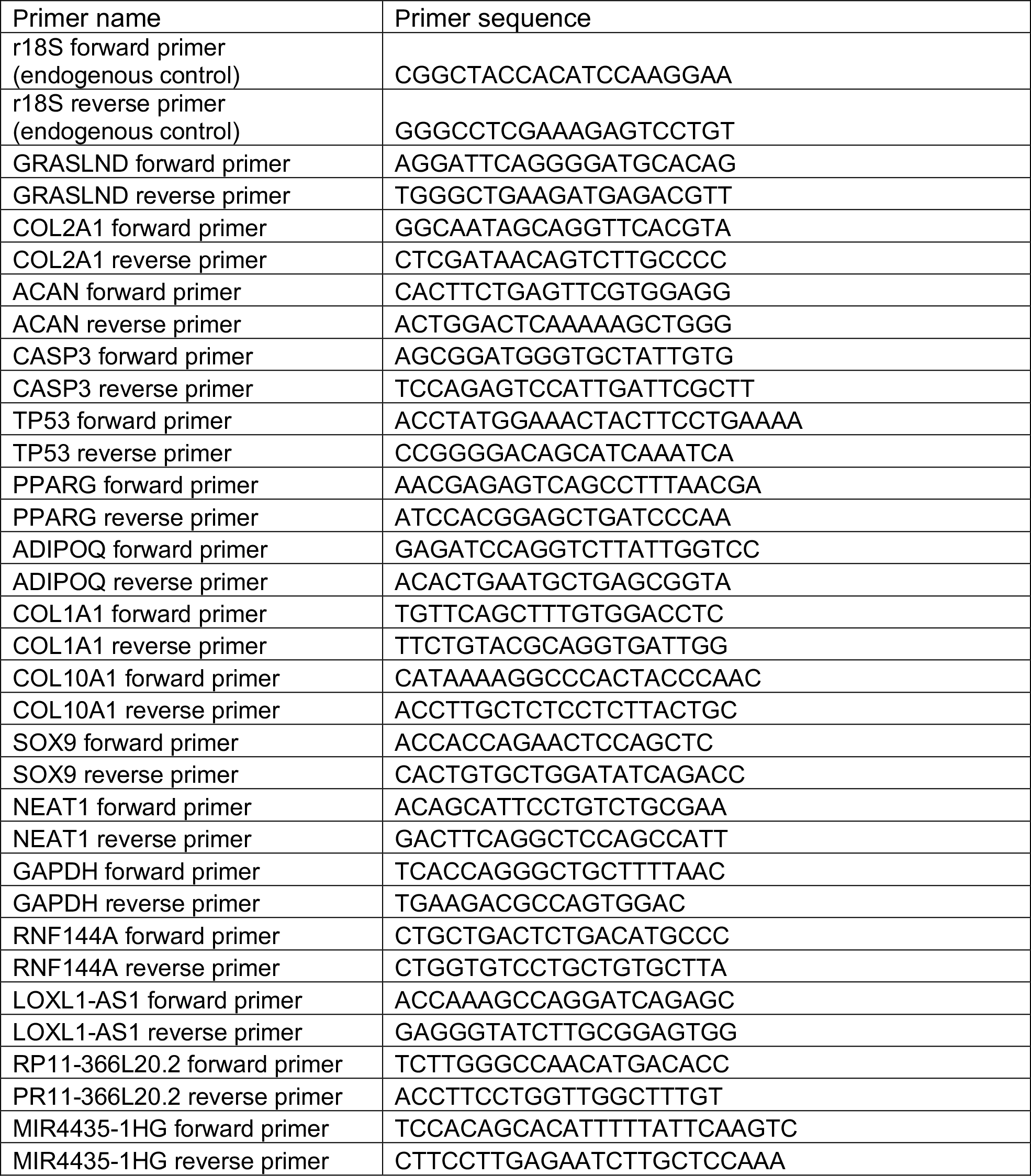
qRT-PCR sequencing primers 5′ – sequence – 3′

### III. Supplemental Materials and Methods

#### Lentivirus production

HEK 293T producer cells were maintained in 293T medium: DMEM-high glucose (Gibco), 10% heat inactivated FBS (Atlas), 1% Penicillin/streptomycin (Gibco). To produce lentivirus for pellet studies, HEK 293T cells were plated at 3.8 × 10^6^ cells per 10 cm dish (Corning) or at 8.3 × 10^6^ cells per 15 cm dish (Falcon) in 293T medium. The following day, cells were co-transfected by calcium phosphate precipitation with the appropriate transfer vector (20 µg for 10 cm dish; 60 µg for 15 cm dish), the second-generation packaging plasmid psPAX2 (Addgene #12260) (15 µg for 10 cm dish; 45 µg for 15 cm dish), and the envelope plasmid pMD2.g (Addgene #12259) (6 µg for 10 cm dish; 18 µg for 15 cm dish). Cells were incubated at 37°C overnight. The following day, fresh medium consisting of DMEM-high glucose (Gibco), 10% heat-inactivated FBS (Atlas), 1% Penicillin/streptomycin (Gibco), 4 mM caffeine (Sigma-Aldrich) was exchanged (12 mL for 10 cm dish; 36 mL for 15 cm dish). Lentivirus was harvested 24 hours post medium change (harvest 1), when fresh medium was exchanged again. 48 hours post medium change, harvest 2 was collected. Harvest 1 and harvest 2 supernatant were pooled, filtered through 0.45 µm cellulose acetate filters (Corning), concentrated, aliquoted and stored at −80°C for future use.

To produce lentivirus for shRNA and gRNA screening, HEK 293T cells were plated at 1.5-2 × 10^6^ cells per well in a 6-well plate in DMEM-high glucose (Gibco), 10% heat inactivated FBS (Atlas). The following day, cells were co-transfected with 2 µg of the appropriate transfer vector, 1.5 µg of the packaging plasmid psPAX2 (Addgene #12260), and 0.6 µg of the envelope plasmid (Addgene #12259) with Lipofectamine 2000 (ThermoFisher) following manufacturer’s protocol. Harvest and storage were performed as described above.

For knockdown experiments, lentivirus was titered by determining the number of antibiotic resistant colonies after puromycin treatment. For overexpression experiments, lentivirus was titered by measuring integrated lentiviral copy number in host DNA with qRT-PCR as previously described (1). Control and tested groups were targeted at similar MOIs.

#### Lentivirus transduction

Cells were plated at 4,500 cells/ cm^2^ for one day and then transduced with appropriate lentivirus in expansion medium supplemented with 4 µg/mL polybrene (Sigma-Aldrich). Twenty-four hours post transduction, cells were rinsed once in phosphate buffered saline (PBS). Cells were cultured with fresh medium exchange every three days until future use.

#### Cytotoxicity assay

Seven days post viral transduction, medium was collected and the amount of lactose dehydrogenase (LDH) was measured as indirect output for cellular toxicity. Assays were performed following manufacturer’s protocol (Promega). Absorbance signal was recorded at 490 nm with the Cytation 5 instrument (BioTek).

#### RNA-seq library preparation

Isolated RNAs were stored at −80°C and submitted to the Genome Technology Access Center at Washington University in St Louis for library preparation and sequencing on a HiSeq 2500 (2 × 101 bp). Libraries were prepared using TruSeq Stranded Total RNA with Ribo-Zero Gold kit (Illumina).

#### RNA pull-down and mass spectrometry

The full sequence of *GRASLND* transcript variant 1 (RefSeq NR_033997.1) was synthesized by Integrated DNA Technologies, Inc, and cloned into the pGEM®-T Easy Vector System (Promega) using the EcoRV site. This served as a template for subsequent in vitro transcription using the Riboprobe® Combination Systems Kit (Promega), with spiked in biotin RNA labeling mix (Roche). Resulted biotinylated sense- and control antisense-transcripts of *GRASLND* were stored at −80°C until further processing.

Cell lysates from day 21 pellets were homogenized in mRIPA buffer (Cell Signaling) and centrifuged at 14,000 rpm for 15 minutes. The protein concentration of cell lysates was measured and adjusted to 2 mg/mL. Five hundred μL of total protein (1mg) were incubated with either 1.5 µg of *GRASLND*-sense or antisense RNA transcripts tagged with biotin-16-UTP overnight (12 hours). Following incubation, the RNA-protein mixtures, and cell lysates (control) were incubated with 100 μL of prewashed streptavidin beads for 3 hours at 4°C (Pierce™ MS-Compatible Magnetic IP Kit, Streptavidin, #90408). The streptavidin beads were then washed five times in 800 µl of ice cold PBS. Beads were eluted twice, each with 30 μL of SDS elution buffer containing 100 mM Tris/HCl pH 8, 4% SDS, 50mM DTT. The elution was either used for mass spectroscopy (Proteomics Core Facility, Washington University School of Medicine), or for Western blot (RayBiotech).

#### Bulk RNA-seq analysis

##### Alignment and Read Assignment

Demultiplexed raw sequencing files were generated by the Genome Technology Access Center at Washington University in St Louis. Reads were processed with trimmomatic-0.36 (2), aligned with STAR-2.6.0 (3), and counted with featureCounts/Subread-1.6.1 (4).

##### Differential Expression Analysis

Downstream differential expression analysis was performed using DESeq2-1.16.1 (5) (abs(log2 fold change) > 1 and adjusted p-values < 0.05).

##### Gene Ontology Analysis

Gene ontology analysis of dysregulated genes was performed with enrichR-1.0 (6, 7).

##### Transcription Factor Identification

Potential transcription factors were identified based on the presence of annotated DNA binding motifs with RcisTarget-1.0.2 (8). Annotation database for the motifs to human transcription factors were previously compiled and can be downloaded at https://resources.aertslab.org/cistarget/. Cis-BP motifs were ranked by normalized enrichment score (NES), and the top five were reported in this paper.

#### Identification of lncRNA candidates

GSE109503 is the dataset that profiles transcriptomic changes of MSC chondrogenesis, composed of six time points (day 0, day 1, day 3, day 7, day 14, day 21) and three biological replicates. Raw sequencing files were downloaded from the GEO Omnibus, and processed as described above. Candidates were first restricted to those differentially expressed per day pair-wise (abs(log2 fold change) > 1 and adjusted p-values < 0.05) and of detectable abundance (TPM > 1 in more than 6 samples across the dataset). lncRNAs whose transcripts were not analyzed for transcript support level (ENSEMBL TSL) were also excluded. For the surviving genes, Pearson correlation analysis was then performed on mean expression per day. Candidates were identified as those with Pearson correlation values > 0.9 to all three investigated markers (ALCAM, VCAM1, ENG for MSC markers; COL2A1, ACAN, COMP for chondrogenic markers; SOX5, SOX6, SOX9 for SOX transcription factors).

GSE69110 depicts the transcriptomic changes of fibroblasts in response to SOX9 expression levels. Raw sequencing files were downloaded from the GEO Omnibus, and processed similarly. Differentially expressed genes between two conditions (SOX9 overexpression versus GFP control) were then identified (abs(log2 fold change) > 1 and adjusted p-values < 0.1). The shortlist of lncRNAs are the intersecting candidates between genes emerging from the above Pearson correlation analysis and dysregulated genes from this dataset.

#### Microarray analysis

Microarray processed data was downloaded from the GEO Omnibus and differential expression analysis was performed with limma-3.34.6 (9).

#### Mass spectrometry analysis

Scaffold-4.8.4 (Proteome Software Inc.) was used to validate MS/MS based peptide and protein identifications. Peptide identifications were accepted if they could be established at greater than 66.0% probability to achieve FDR less than 1.0% by the Scaffold Local FDR algorithm. Protein identification was accepted if they could be established at a greater than 95.0% probability and contained at least one identified peptide. Protein probabilities were assigned by the Protein Prophet algorithm (10). Proteins that contain similar peptides and could not be differentiated based on MS/MS analysis alone were grouped to satisfy the principles of parsimony. To identify differentially bound proteins, one tailed t-test was performed on sense samples compared to naked beads, and sense samples compared to antisense samples.

## References

1. Sophia Fox AJ, Bedi A, & Rodeo SA (2009) The basic science of articular cartilage: structure, composition, and function. Sports Health 1(6):461–468.

2. Huynh NPT, et al. (2018) Genetic Engineering of Mesenchymal Stem Cells for Differential Matrix Deposition on 3D Woven Scaffolds. Tissue Eng Part A 24(19-20):1531–1544.

3. Glass KA, et al. (2014) Tissue-engineered cartilage with inducible and tunable immunomodulatory properties. Biomaterials 35(22):5921–5931.

4. Brunger JM, Zutshi A, Willard VP, Gersbach CA, & Guilak F (2017) CRISPR/Cas9 Editing of Murine Induced Pluripotent Stem Cells for Engineering Inflammation-Resistant Tissues. Arthritis Rheumatol 69(5):1111–1121.

5. Brunger JM, Zutshi A, Willard VP, Gersbach CA, & Guilak F (2017) Genome Engineering of Stem Cells for Autonomously Regulated, Closed-Loop Delivery of Biologic Drugs. Stem Cell Reports 8(5):1202–1213.

6. Brunger JM, et al. (2014) Scaffold-mediated lentiviral transduction for functional tissue engineering of cartilage. Proc Natl Acad Sci U S A 111(9):E798–806.

7. Adkar SS, et al. (2017) Genome Engineering for Personalized Arthritis Therapeutics. Trends Mol Med 23(10):917–931.

8. Gimble J & Guilak F (2003) Adipose-derived adult stem cells: isolation, characterization, and differentiation potential. Cytotherapy 5(5):362–369.

9. Erickson GR, et al. (2002) Chondrogenic potential of adipose tissue-derived stromal cells in vitro and in vivo. Biochem Biophys Res Commun 290(2):763–769.

10. Awad HA, Wickham MQ, Leddy HA, Gimble JM, & Guilak F (2004) Chondrogenic differentiation of adipose-derived adult stem cells in agarose, alginate, and gelatin scaffolds. Biomaterials 25(16):3211–3222.

11. Caplan AI (1991) Mesenchymal stem cells. J Orthop Res 9(5):641–650.

12. Mackay AM, et al. (1998) Chondrogenic differentiation of cultured human mesenchymal stem cells from marrow. Tissue Eng 4(4):415–428.

13. Johnstone B, Hering TM, Caplan AI, Goldberg VM, & Yoo JU (1998) In vitro chondrogenesis of bone marrow-derived mesenchymal progenitor cells. Exp Cell Res 238(1):265–272.

14. Lander ES, et al. (2001) Initial sequencing and analysis of the human genome. Nature 409(6822):860–921.

15. Guttman M, et al. (2009) Chromatin signature reveals over a thousand highly conserved large non-coding RNAs in mammals. Nature 458(7235):223–227.

16. Guttman M, et al. (2011) lincRNAs act in the circuitry controlling pluripotency and differentiation. Nature 477(7364):295–300.

17. Lee JT & Bartolomei MS (2013) X-inactivation, imprinting, and long noncoding RNAs in health and disease. Cell 152(6):1308–1323.

18. Ng SY, Johnson R, & Stanton LW (2012) Human long non-coding RNAs promote pluripotency and neuronal differentiation by association with chromatin modifiers and transcription factors. EMBO J 31(3):522–533.

19. Briggs JA, Wolvetang EJ, Mattick JS, Rinn JL, & Barry G (2015) Mechanisms of Long Non-coding RNAs in Mammalian Nervous System Development, Plasticity, Disease, and Evolution. Neuron 88(5):861–877.

20. Carlson HL, et al. (2015) LncRNA-HIT Functions as an Epigenetic Regulator of Chondrogenesis through Its Recruitment of p100/CBP Complexes. PLoS Genet 11(12):e1005680.

21. Barter MJ, et al. (2017) The long non-coding RNA ROCR contributes to SOX9 expression and chondrogenic differentiation of human mesenchymal stem cells. Development 144(24):4510–4521.

22. Huynh NP, Anderson BA, Guilak F, & McAlinden A (2017) Emerging roles for long noncoding RNAs in skeletal biology and disease. Connect Tissue Res 58(1):116–141.

23. Dieudonne FX, et al. (2013) Promotion of osteoblast differentiation in mesenchymal cells through Cbl-mediated control of STAT5 activity. Stem Cells 31(7):1340–1349.

24. Rostovskaya M, et al. (2018) Clonal Analysis Delineates Transcriptional Programs of Osteogenic and Adipogenic Lineages of Adult Mouse Skeletal Progenitors. Stem Cell Reports 11(1):212–227.

25. Takayanagi H, et al. (2002) RANKL maintains bone homeostasis through c-Fos-dependent induction of interferon-beta. Nature 416(6882):744–749.

26. Takayanagi H, Kim S, & Taniguchi T (2002) Signaling crosstalk between RANKL and interferons in osteoclast differentiation. Arthritis Res 4 Suppl 3:S227–232.

27. Li J (2013) JAK-STAT and bone metabolism. JAKSTAT 2(3):e23930.

28. Sahni M, et al. (1999) FGF signaling inhibits chondrocyte proliferation and regulates bone development through the STAT-1 pathway. Genes Dev 13(11):1361–1366.

29. Jang YN & Baik EJ (2013) JAK-STAT pathway and myogenic differentiation. JAKSTAT 2(2):e23282.

30. Xiao L, et al. (2004) Stat1 controls postnatal bone formation by regulating fibroblast growth factor signaling in osteoblasts. J Biol Chem 279(26):27743–27752.

31. Sahni M, Raz R, Coffin JD, Levy D, & Basilico C (2001) STAT1 mediates the increased apoptosis and reduced chondrocyte proliferation in mice overexpressing FGF2. Development 128(11):2119–2129.

32. Brierley MM & Fish EN (2002) Review: IFN-alpha/beta receptor interactions to biologic outcomes: understanding the circuitry. J Interferon Cytokine Res 22(8):835–845.

33. Hertzog PJ, Hwang SY, & Kola I (1994) Role of interferons in the regulation of cell proliferation, differentiation, and development. Mol Reprod Dev 39(2):226–232.

34. Hu X & Ivashkiv LB (2009) Cross-regulation of signaling pathways by interferon-gamma: implications for immune responses and autoimmune diseases. Immunity 31(4):539–550.

35. Green DS, Young HA, & Valencia JC (2017) Current prospects of type II interferon gamma signaling and autoimmunity. J Biol Chem 292(34):13925–13933.

36. Huynh NPT, Zhang B, & Guilak F (2018) High-depth transcriptomic profiling reveals the temporal gene signature of human mesenchymal stem cells during chondrogenesis. FASEB J:fj201800534R.

37. Ohba S, He X, Hojo H, & McMahon AP (2015) Distinct Transcriptional Programs Underlie Sox9 Regulation of the Mammalian Chondrocyte. Cell Rep 12(2):229–243.

38. Clemson CM, et al. (2009) An architectural role for a nuclear noncoding RNA: NEAT1 RNA is essential for the structure of paraspeckles. Mol Cell 33(6):717–726.

39. Sasaki YT, Ideue T, Sano M, Mituyama T, & Hirose T (2009) MENepsilon/beta noncoding RNAs are essential for structural integrity of nuclear paraspeckles. Proc Natl Acad Sci U S A 106(8):2525–2530.

40. Rhesus Macaque Genome S, et al. (2007) Evolutionary and biomedical insights from the rhesus macaque genome. Science 316(5822):222–234.

41. Derrien T, et al. (2012) The GENCODE v7 catalog of human long noncoding RNAs: analysis of their gene structure, evolution, and expression. Genome Res 22(9):1775–1789.

42. Necsulea A, et al. (2014) The evolution of lncRNA repertoires and expression patterns in tetrapods. Nature 505(7485):635–640.

43. Black JB, et al. (2016) Targeted Epigenetic Remodeling of Endogenous Loci by CRISPR/Cas9-Based Transcriptional Activators Directly Converts Fibroblasts to Neuronal Cells. Cell Stem Cell 19(3):406–414.

44. Perez-Pinera P, et al. (2013) RNA-guided gene activation by CRISPR-Cas9-based transcription factors. Nat Methods 10(10):973–976.

45. Ramos YF, et al. (2014) Genes involved in the osteoarthritis process identified through genome wide expression analysis in articular cartilage; the RAAK study. PLoS One 9(7):e103056.

46. Dunn SL, et al. (2016) Gene expression changes in damaged osteoarthritic cartilage identify a signature of non-chondrogenic and mechanical responses. Osteoarthritis Cartilage 24(8):1431–1440.

47. Cheng M, Nguyen MH, Fantuzzi G, & Koh TJ (2008) Endogenous interferon-gamma is required for efficient skeletal muscle regeneration. Am J Physiol Cell Physiol 294(5):C1183–1191.

48. Londhe P & Davie JK (2011) Gamma interferon modulates myogenesis through the major histocompatibility complex class II transactivator, CIITA. Mol Cell Biol 31(14):2854–2866.

49. Ishida Y, Kondo T, Takayasu T, Iwakura Y, & Mukaida N (2004) The essential involvement of cross-talk between IFN-gamma and TGF-beta in the skin wound-healing process. J Immunol 172(3):1848–1855.

50. Yufit T, Vining V, Wang L, Brown RR, & Varga J (1995) Inhibition of type I collagen mRNA expression independent of tryptophan depletion in interferon-gamma-treated human dermal fibroblasts. J Invest Dermatol 105(3):388–393.

51. Harrop AR, et al. (1995) Regulation of collagen synthesis and mRNA expression in normal and hypertrophic scar fibroblasts in vitro by interferon-gamma. J Surg Res 58(5):471–477.

52. Granstein RD, Flotte TJ, & Amento EP (1990) Interferons and collagen production. J Invest Dermatol 95(6 Suppl):75S–80S.

53. Amento EP, Bhan AK, McCullagh KG, & Krane SM (1985) Influences of gamma interferon on synovial fibroblast-like cells. Ia induction and inhibition of collagen synthesis. J Clin Invest 76(2):837–848.

54. Wang S, et al. (2006) The catalytic activity of the eukaryotic initiation factor-2alpha kinase PKR is required to negatively regulate Stat1 and Stat3 via activation of the T-cell protein-tyrosine phosphatase. J Biol Chem 281(14):9439–9449.

55. Wong AH, et al. (1997) Physical association between STAT1 and the interferon-inducible protein kinase PKR and implications for interferon and double-stranded RNA signaling pathways. EMBO J 16(6):1291–1304.

56. Osman F, Jarrous N, Ben-Asouli Y, & Kaempfer R (1999) A cis-acting element in the 3′-untranslated region of human TNF-alpha mRNA renders splicing dependent on the activation of protein kinase PKR. Genes Dev 13(24):3280–3293.

57. Ben-Asouli Y, Banai Y, Pel-Or Y, Shir A, & Kaempfer R (2002) Human interferon-gamma mRNA autoregulates its translation through a pseudoknot that activates the interferon-inducible protein kinase PKR. Cell 108(2):221–232.

58. Cohen-Chalamish S, et al. (2009) Dynamic refolding of IFN-gamma mRNA enables it to function as PKR activator and translation template. Nat Chem Biol 5(12):896–903.

59. Nallagatla SR, et al. (2007) 5′-triphosphate-dependent activation of PKR by RNAs with short stem-loops. Science 318(5855):1455–1458.

60. Mayo CB & Cole JL (2017) Interaction of PKR with single-stranded RNA. Sci Rep 7(1):3335.

61. Boissier MC, et al. (1995) Biphasic effect of interferon-gamma in murine collagen-induced arthritis. Eur J Immunol 25(5):1184–1190.

62. Cooper SM, Sriram S, & Ranges GE (1988) Suppression of murine collagen-induced arthritis with monoclonal anti-Ia antibodies and augmentation with IFN-gamma. J Immunol 141(6):1958–1962.

63. Westacott CI, et al. (1990) Synovial fluid concentration of five different cytokines in rheumatic diseases. Ann Rheum Dis 49(9):676–681.

64. Kahle P, et al. (1992) Determination of cytokines in synovial fluids: correlation with diagnosis and histomorphological characteristics of synovial tissue. Ann Rheum Dis 51(6):731–734.

65. Bester AC, et al. (2018) An Integrated Genome-wide CRISPRa Approach to Functionalize lncRNAs in Drug Resistance. Cell 173(3):649–664 e620.

66. Liu SJ, et al. (2017) CRISPRi-based genome-scale identification of functional long noncoding RNA loci in human cells. Science 355(6320).

67. Hagmann S, et al. (2013) FGF-2 addition during expansion of human bone marrow-derived stromal cells alters MSC surface marker distribution and chondrogenic differentiation potential. Cell Prolif 46(4):396–407.

68. Moffat J, et al. (2006) A lentiviral RNAi library for human and mouse genes applied to an arrayed viral high-content screen. Cell 124(6):1283–1298.

69. Diekman BO, et al. (2015) Knockdown of the cell cycle inhibitor p21 enhances cartilage formation by induced pluripotent stem cells. Tissue Eng Part A 21(7-8):1261–1274.

70. Barde I, et al. (2006) Efficient control of gene expression in the hematopoietic system using a single Tet-on inducible lentiviral vector. Mol Ther 13(2):382–390.

71. Kent WJ, et al. (2002) The human genome browser at UCSC. Genome Res 12(6):996–1006.

72. Doench JG, et al. (2016) Optimized sgRNA design to maximize activity and minimize off-target effects of CRISPR-Cas9. Nat Biotechnol 34(2):184–191.

73. Haeussler M, et al. (2016) Evaluation of off-target and on-target scoring algorithms and integration into the guide RNA selection tool CRISPOR. Genome Biol 17(1):148.

74. Kabadi AM, Ousterout DG, Hilton IB, & Gersbach CA (2014) Multiplex CRISPR/Cas9-based genome engineering from a single lentiviral vector. Nucleic Acids Res 42(19):e147.

75. Sastry L, Johnson T, Hobson MJ, Smucker B, & Cornetta K (2002) Titering lentiviral vectors: comparison of DNA, RNA and marker expression methods. Gene Ther 9(17):1155–1162.

76. Farndale RW, Buttle DJ, & Barrett AJ (1986) Improved quantitation and discrimination of sulphated glycosaminoglycans by use of dimethylmethylene blue. Biochim Biophys Acta 883(2):173–177.

77. Estes BT, Diekman BO, Gimble JM, & Guilak F (2010) Isolation of adipose-derived stem cells and their induction to a chondrogenic phenotype. Nat Protoc 5(7):1294–1311.

78. Team RC (2018) R: A language and environment for statistical computing. R Foundation for Statistical Computing Vienna, Austria.

79. Weirauch MT, et al. (2014) Determination and inference of eukaryotic transcription factor sequence specificity. Cell 158(6):1431–1443.

## References

1. Sastry L, Johnson T, Hobson MJ, Smucker B, & Cornetta K (2002) Titering lentiviral vectors: comparison of DNA, RNA and marker expression methods. Gene Ther 9(17):1155–1162.

2. Bolger AM, Lohse M, & Usadel B (2014) Trimmomatic: a flexible trimmer for Illumina sequence data. Bioinformatics 30(15):2114–2120.

3. Dobin A, et al. (2013) STAR: ultrafast universal RNA-seq aligner. Bioinformatics 29(1):15–21.

4. Liao Y, Smyth GK, & Shi W (2014) featureCounts: an efficient general purpose program for assigning sequence reads to genomic features. Bioinformatics 30(7):923–930.

5. Love MI, Huber W, & Anders S (2014) Moderated estimation of fold change and dispersion for RNA-seq data with DESeq2. Genome Biol 15(12):550.

6. Chen EY, et al. (2013) Enrichr: interactive and collaborative HTML5 gene list enrichment analysis tool. BMC Bioinformatics 14:128.

7. Kuleshov MV, et al. (2016) Enrichr: a comprehensive gene set enrichment analysis web server 2016 update. Nucleic Acids Res 44(W1):W90–97.

8. Aibar S, et al. (2017) SCENIC: single-cell regulatory network inference and clustering.Nat Methods 14(11):1083–1086.

9. Ritchie ME, et al. (2015) limma powers differential expression analyses for RNA-sequencing and microarray studies. Nucleic Acids Res 43(7):e47.

10. Nesvizhskii AI, Keller A, Kolker E, & Aebersold R (2003) A statistical model for identifying proteins by tandem mass spectrometry. Anal Chem 75(17):4646–4658.

